# Characterization of a novel fatty acid-modifying pathway toward the biosynthesis of tambjamine BE-18591 in *Streptomyces*

**DOI:** 10.1101/2025.08.11.669332

**Authors:** Neil L. Grenade, Yan Feng, Emily H. Perrino, Avena C. Ross, Graeme W. Howe

**Affiliations:** Department of Chemistry, Queen’s University, Kingston, ON, K7L 3N6, Canada. Tambjamine, Prodiginine, Enzymology, Natural Products, Biosynthesis

## Abstract

Tambjamines are a class of bacterial bipyrrolic natural products with potent biological activity. Recently, the first Actinomycete biosynthetic gene cluster (BGC) responsible for the production of a tambjamine (BE-18591) was discovered in *Streptomyces albus*, and bioinformatic analysis suggested the alkylamine tail component is constructed using disparate biosynthetic logic than employed by Proteobacteria to assemble structurally similar tambjamines YP1 and MYP1. Here, we report the experimental characterization of four novel streptomycete proteins and demonstrate that these enable the unique, late-stage assembly of the alkylamine component of BE-18951. Specifically, a fatty acyl-carrier protein (TabQ) is loaded with a 12-carbon acyl chain, selected for, in part, through the action of an editing type II thioesterase (TabJ). The resulting C_12_-TabQ adduct is then processed to an aldehyde by a novel acyl-ACP reductase (TabE) that harbors none of the telltale amino acid signatures that typically identify these proteins. The resulting aldehyde is finally converted to the amine by an *ω*-transaminase (TabA) that demonstrates some degree of promiscuity. These four proteins encoded by the BE-18591 BGC in *Streptomyces albus* enable the assembly of the fatty amine that is ultimately incorporated into the tambjamine. Our findings highlight the disparate chemical logic employed by Proteo- and Actinobacteria for the biosynthesis of the alkylamine components of tambjamine natural products.

## INTRODUCTION

Tambjamines and prodiginines are related classes of bipyrrolic natural products (NPs) with therapeutic potential as antimicrobial,^1,2^ antineoplastic^3^ and immunomodulatory^4^ drugs. Both classes contain a conserved core derived from 4’-methoxy*-*2,2’-bipyrrole-5’-carbaldehyde (MBC; **1**; Scheme 1a) that is either coupled to a third pyrrole (prodiginines) or an alkylamine (tambjamines). Known producers of these NPs include both Proteobacteria and Actinobacteria, although comparatively little is known about the biosynthesis of tambjamines relative to prodiginines.^5^ Recently, we described the second tambjamine biosynthetic gene cluster (BGC) and the first from an actinobacterium, revealing significant differences between the biosynthetic logic employed by Actino- and Proteobacteria to assemble the alkylamine tail of these NPs (Scheme 1).^6^ Tambjamine YP1 (**2**; Scheme 1a) is encoded by the *tam* BGC in the proteobacteria *Pseudoalteromonas tunicata* and *Pseudoalteromonas citrea,* while tambjamine BE-18591 (**3**; Scheme 1b) is encoded by the *tab* BGC in the actinobacterium *Streptomyces albus.* Tambjamines **2** and **3** are nearly identical in structure, differing only by a single degree of unsaturation within the alkylamine tail. Initial analysis of the corresponding BGCs suggested that while the MBC core is likely produced the same way in both systems, the construction of the fatty acyl precursor of the alkylamine tails was predicted to differ significantly. While the tail in tambjamine **2** was postulated to be derived from the existing pool of fatty acids derived from primary metabolism, the saturated tail in the actinobacterial tambjamine **3** was proposed to be assembled by several dedicated enzymes encoded within the *tab* BGC.^6,7^ Specifically, the Tam biosynthetic pathway was proposed to use three bifunctional enzymes to convert the 12-carbon primary metabolite dodecanoic acid (**4**; Scheme 1a) to the corresponding unsaturated fatty amine *cis*-dodec-3-en-1-amine (DDEA; **5**; Scheme 1a). Campopiano has shown that TamA performs the adenosine triphosphate (ATP)- dependent adenylation and thiolation of **4**,^7^ and that TamH catalyzes the NADH-dependent reductive offloading of the acyl chain from TamA and transamination of the resulting aldehyde to the unsaturated amine tail (**5**; Scheme 1a).^8^ When these authors evaluated the activities of TamA and TamH with substrates with varying tail lengths,^7,8^ both enzymes gave maximal conversions with dodecanoic acid-derived substrates. Further, while TamH catalyzed the production of dodecylamine (**6**; Scheme 1a) from a saturated C_12_-TamA adduct,^8^ no activity was observed with the corresponding acyl-coenzyme A (CoA) derivative, suggesting that interactions with the loaded acyl carrier protein (ACP) are required for the thioreductase and/or transaminase activity of TamH. While the production of **5** is hypothesized to require oxidation and isomerization of a TamA-tethered substrate by the flavin dinucleotide (FAD)-dependent TamT to generate the *Z*-alkene in **5** (Scheme 1a), this step has not yet been experimentally verified.

At the outset of this work, comparatively little was known about the biosynthetic machinery involved in assembling the alkylamine tail of BE-18591. Functional annotation of the *tab* BGC identified *tabA, tabE, tabJ, tabP, tabQ,* and *tabR* as genes with possible relevance to dodecylamine biosynthesis in *Streptomyces*, but the corresponding enzymes had not been characterized experimentally. While *tabP* and *tabR* putatively encode dedicated β-ketoacyl synthases for fatty acid synthesis,^6^ the roles of several remaining genes were postulated to encode tailoring enzymes to convert the ACP-harbored alkyl chain to dodecylamine (**6**; Scheme 1b). Specifically, TabJ was predicted to be a thioesterase responsible for catalyzing the hydrolysis of the mature twelve-carbon acyl chain off of TabQ, the putative acyl carrier protein (ACP). TabE was annotated as an NAD(P)H-dependent aldehyde dehydrogenase (ALDH), and while ALDHs nominally oxidize aldehydes to carboxylates, TabE was hypothesized to instead catalyze the thermodynamically uphill reduction of dodecanoic acid to the corresponding aldehyde. Finally, this aldehyde was predicted to be converted to the alkylamine (**6**) by the action of the pyridoxal-5′-phosphate (PLP)-dependent transaminase TabA.^6^

Here, we describe the heterologous overexpression, purification, and biochemical characterization of the four proteins involved in the production of **6** in *Streptomyces albus*. *In vitro* assays were used to evaluate the proposed biosynthesis of **6**, with results reported here requiring a revised scheme wherein TabJ is not responsible for productive acyl chain offloading from TabQ. Instead, TabE catalyzes the reductive offloading of the TabQ-loaded thioester to the corresponding aldehyde, which is subsequently converted to **6** by the action of the TabA transaminase. The revised scheme invokes TabJ as a type II thioesterase responsible for “editing” the TabQ-loaded acyl chain by hydrolyzing aberrant acyl groups, a function that is supported by the promiscuous hydrolytic activity reported in this work. Given the proposed convergent evolution of the machinery responsible for alkylamine tail biosynthesis in proteo- and actinobacterial tambjamines, the revised biosynthetic scheme brought about by this work has important implications for understanding how bipyrrolic NPs emerged in these different phyla.

## RESULTS AND DISCUSSION

### TabQ is a discrete type II PKS-like fatty acyl-carrier protein

Based on bioinformatic analysis, TabQ (NCBI accession: WP_016467649.1) was annotated as an ACP. Alignment of the TabQ sequence with other functionally characterized ACPs suggested that a highly conserved serine residue (S35; Figure S1) is responsible for covalently anchoring the 4′-phosphopantetheine (Ppant) arm to the protein (Figure 1a). Additionally, a putative “recognition” helix motif (residues 36-49) was identified in the predicted structure of TabQ that could facilitate the interaction of acyl chain-modifying enzymes with this stand-alone type II polyketide synthase (PKS)-like ACP.^9^ Despite putatively performing similar functions, facilitating modification of tethered acyl substrates, TabQ shares only 19.1% sequence identity with the ACP domain of TamA (Figure S1). This is consistent with previous demonstrations of the sequence variability within the ACP-like superfamily, which suggested the existence of 16 distinct subfamilies of ACP.^10^ The ACP domain of TamA has highest sequence alignment with the ACP12 family, which consists of multidomain enzymes like enterobactin synthase, a type I PKS. The presence of a fused adenylation domain in TamA alongside the ACP domain conforms with the multidomain classification. In contrast, TabQ aligns most closely with the ACP5 family, which is composed of standalone ACPs associated with type II PKSs.^10^ In an alignment of representative members of subclasses ACP5 and ACP12, the subclasses were reported to share approximately 15.8% sequence identity with each other.^10^ The comparable %ID obtained by the direct alignment of TabQ and TamA is consistent with the putative assignment of these ACPs to subclasses 5 and 12, respectively.

**Figure 1.**
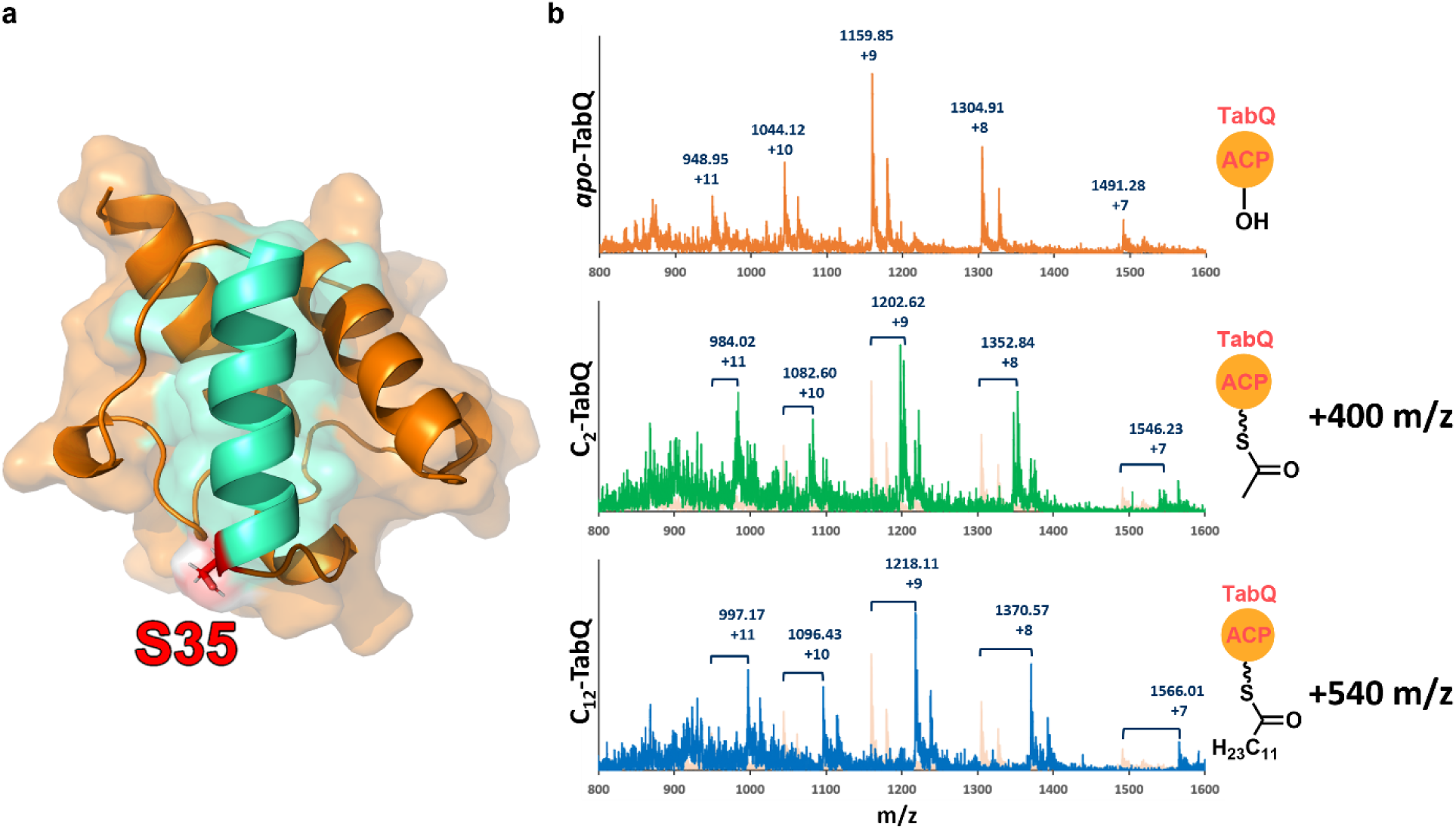
**a**) AlphaFold2^11^ predicted structure of the discrete acyl carrier protein, apo-TabQ. Ser35 (shown in red) is functionalized with an acyl or non-acyl bound-4′-phosphopanteheine (Ppant) group to form acyl- or *holo*-TabQ, respectively. The “recognition helix” (residues 36-49) putatively recognized by downstream acyl chain processing enzymes is shown in turquoise. **b**) Mass spectra showing m/z increases (charge states +7 to +11) corresponding to the loading of C_2_- and C_12_-Ppant onto *apo*-TabQ. Spectrum from *apo*-TabQ is underlayed in C_2_-TabQ and C_12_-TabQ spectra.

Prior to *in vitro* assays with TabQ, the *apo* protein was converted to the *holo* form using recombinant Sfp from *Bacillus subtilis* as a phospantetheinyl transferase (PPTase) and an acyl-CoA as an acyl-Ppant donor. The successful loading of C_2_- and C_12_-Ppant groups onto TabQ was confirmed by LC-MS (Figure 1b). As the alkyl tail found in **3** is 12 carbons long, C_12_-TabQ was initially presumed to be the preferred substrate for downstream enzyme, TabJ. As such, assays with both TabJ and TabE were conducted with C_12_-TabQ to evaluate the next step in the biosynthesis of **6**.

### TabJ is a type II thioesterase with broad substrate selectivity

Initially, TabJ (NCBI accession: WP_106963665.1) had been invoked as a thioesterase responsible for hydrolyzing the appropriately formed TabQ-loaded acyl chain.^6^ However, the presence of a GxSMG consensus motif centered on the putative catalytic Ser76 suggests that TabJ is an “editing” type II thioesterase (TEII) that could instead catalyze the removal of malformed TabQ-loaded acyl chains (Figure S2).**^Error! Reference source not found.^**^,13^ The predicted structure of TabJ includes an “open” substrate channel leading to a serine-histidine-aspartate catalytic triad at the interface of the enzyme core and lid regions (Figures 2a and 2b). Similar structural motifs have been reported for other TEIIs including RedJ (51.7% ID), the TEII from the undecylprodigiosin _biosynthetic pathway._**_Error! Reference source not found._**,_1_^5^

**Figure 2.**
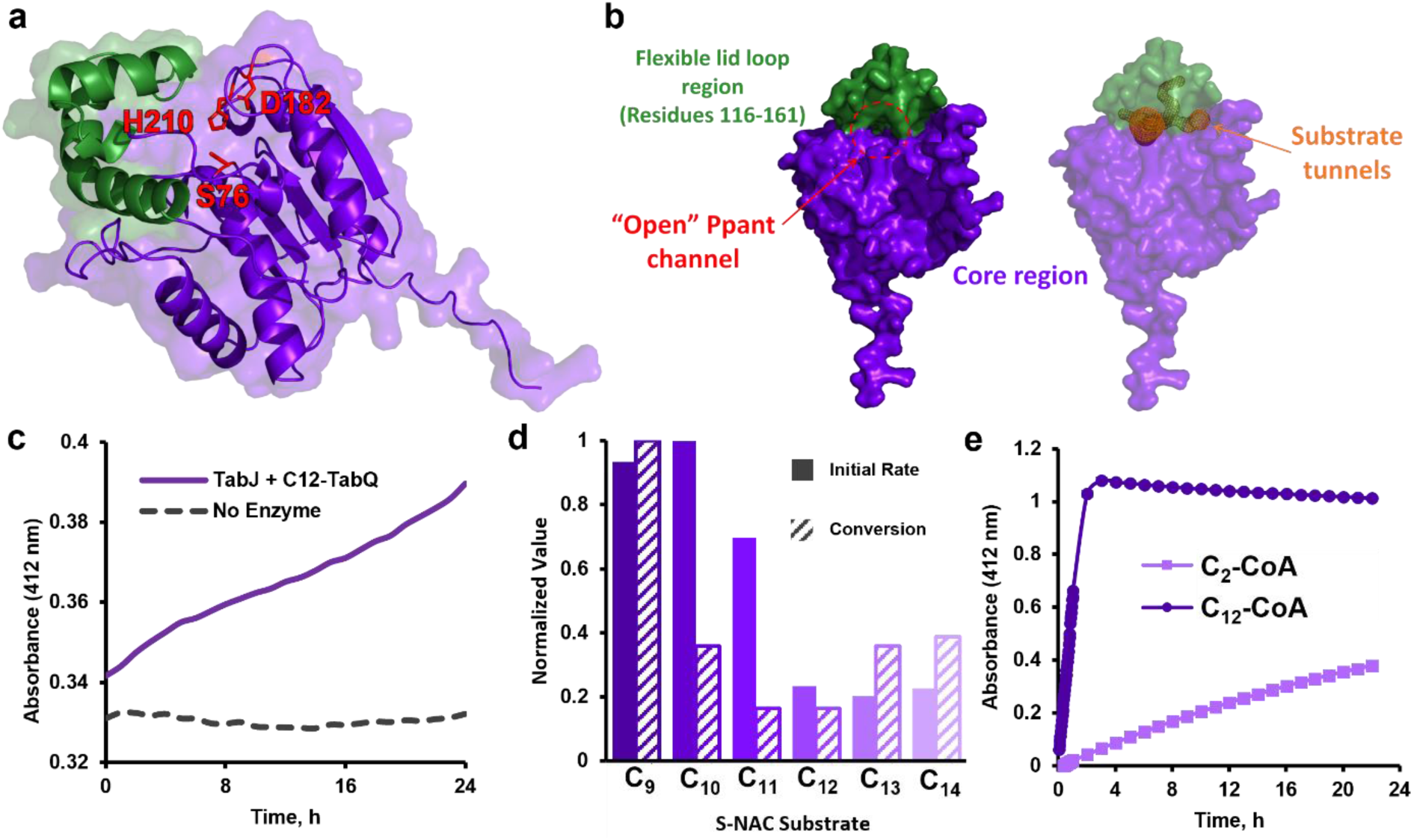
**a)** AlphaFold2^11^ predicted structure of TabJ. Residues that comprise the putative catalytic triad are highlighted in red. The structure of TabJ is divided into the core (purple) and lid (green) **b**) The “open” Ppant channel at the interface of the flexible lid and core regions align with the computationally predicted substrate tunnels (orange). The “open” Ppant channel at the interface of the flexible lid and core regions aligns with the Caver 3.0-predicted substrate tunnels (orange). **d)** *In vitro* assays demonstrate the hydrolytic activity of TabJ against acyl coenzyme A thioesters. **e)** SNAC thioesters with shorter alkyl chains are hydrolyzed more efficiently by TabJ *in vitro*. **e)** TabJ catalyzes the hydrolysis of the C_12_-TabQ thioester *in vitro*.

TEIIs typically exhibit “editing” activity by cleaving aberrant ACP-loaded acyl chains.^15^ When the activity of TabJ toward the C_12_-TabQ thioester was monitored using Ellman’s assay (Scheme S1),^16^ absorbance changes at 412 nm indicated that TabJ catalyzed the hydrolysis of this substrate against a no-enzyme control with modest efficiency (Figure 2c). Specifically, with an extinction coefficient of ε_412_ = 13,600 M^-1^cm^-1^ for the TNB anion (Scheme S1), the absorbance increase observed after co-incubation of C_12_-TabQ with TabJ suggests that only 4.4 µM of free thiol was released over the course of 24 hours. This poor (but unambiguous) activity suggested that rather than serving to process the C-12 pathway intermediate, TabJ might instead be a TEII that preferentially hydrolyzes acyl groups of undesired lengths off of TabQ. Difficulties associated with accessing acyl-loaded TabQ substrates made explicit evaluation of this editing hypothesis infeasable. Instead, the ability of TabJ to cleave thioester analogues with alkyl tails of various lengths was investigated. *S*-*N*-acetylcysteamine (SNAC) thioesters of fatty acids with tails ranging from nine to fourteen carbons-long were synthesized to mimic the thioester portion of the Ppant arm with the putative acyl-ACP substrates attached (Scheme S2). All SNACs evaluated as potential substrates for TabJ were hydrolyzed by the enzyme, with compounds with longer acyl chains generally being processed less efficiently (Figure 2d). Notably, while the C_12_-SNAC was also hydrolyzed by TabJ, the relative rates of conversion and the degree of hydrolysis following the assay were minimal when compared to the results obtained with the other SNACs. This result is consistent with the activity of TabJ as a TEII that effectively selects for acyl chains of the correct length for incorporation into the final NP by hydrolyzing acyl chains of the incorrect length.

C_2_-CoA and C_12_-CoA were also evaluated as substrates for TabJ, and while this assay clearly indicated that TabJcatalyzed the hydrolysis of both C_2_-CoA or C_12_-CoA, the C_12_-CoA substrate gave rise to significantly higher initial rates, as well as higher yields of product following the 22-hour assay (Figure 2e). This seemed to contradict the TEII function suggested by results obtained with the SNACs, since the substrate with the “correct” tail length C_12_-CoA was preferentially hydrolyzed over the C_2_-CoA. This apparent discrepancy might arise from the necessity for TabJ to be less active towards very short chain acyl groups during the initial construction of the growing acyl chain. It is thus plausible that TabJ has the greatest activity toward acyl thioesters formed after the initial extension (>4 carbons in length) given the poor activity toward C_2_-CoA. We also note that the activity of TabJ toward acyl-CoA substrates demonstrates that acyl thioesters need not be anchored to TabQ to be hydrolyzed by TabJ and suggests that TabJ is somewhat promiscuous.

While direct comparisons of the rates of hydrolysis of acyl-CoA thioesters and the TabQ-loaded thioesters must be treated with caution, we note the large discrepancy in apparent rates of hydrolysis with C_12_-CoA (Figure 2e) and C_12_-TabQ (Figure 2c). The absorbance change observed when C_12_-CoA was incubated with TabJ indicated that approximately 73.5 µM free thiol was released in less than two hours. While the concentration of the loaded TabQ was likely much lower than the C_12_-CoA employed in these assays, the slow hydrolysis of the C_12_-TabQ seems to represent an inefficient, basal activity toward the thioester substrate with the 12-carbon tail required for the condensation step toward **2**. It is plausible that this basal activity permits TabJ to exert some control on titres of **2**,^14^ or perhaps to select the correct starter unit (malonyl-CoA) for the elongation of the fatty acyl-TabQ intermediate.^17^ As such, we postulate that the results obtained with TabJ are most consistent with this enzyme being a TEII responsible for the preferential elimination of non-productive thioesters of incorrect length from the pool of potential substrates for downstream enzymes in the BE-18591 biosynthetic pathway. Lastly, we note that no functional orthologues of TabJ are present in the Tam pathway, but C_12_ chain selectivity was previously reported in the acyl chain processing enzymes TamA and TamH, which may substitute the need for an editing TEII in the biosynthesis of **5**.^7,8^

### TabE is a novel fatty acyl-ACP reductase

Based on sequence analysis, TabE (NCBI accession: WP_031174883.1) is annotated as an aldehyde dehydrogenase (ALDH). Sequence alignment with authentic ALDHs revealed the conservation of an active-site cysteine nucleophile, a glutamate base, and oxyanion-stabilizing asparagine, consistent with the annotation of TabE as an ALDH (Figure 3a and Figure S3).^18^ Within the context of the biosynthesis of BE-18591, TabE was proposed to reduce dodecanoic acid, released by the action of TabJ on C_12_-TabQ, to the free aldehyde (Scheme 1a).^6^ The carboxylates of free fatty acids are exceptionally poor electrophiles, such that Nature generally drives the kinetically and thermodynamically unfavourable reductions of fatty acids forward by first activating the acyl group by adenylation or thiolation. Relevant examples of this activity are found in NADPH- and ATP-dependent carboxylic acid reductases (CARs) and acyl-CoA reductases that accept carboxylates that have been derivatized to thioesters.^19,20,21,22^ *In silico* analysis of the TabE sequence using IntroProScan^23^ did not identify an ATP-grasp domain in this protein, nor did this analysis indicate that TabE was a member of any recognized acyl-CoA reductase family. Furthermore, sequence similarity networks (SSNs),^24,25^ MIBiG,^26^ and AntiSMASH^27^ revealed no functionally characterized NP biosynthetic orthologues of TabE, and the AntiSMASH database contained no BGCs suggested to utilize an ALDH-like enzyme to reduce a free fatty acid in a manner analogous to the proposed transformation in the Tab pathway (Scheme 1a).

**Figure 3.**
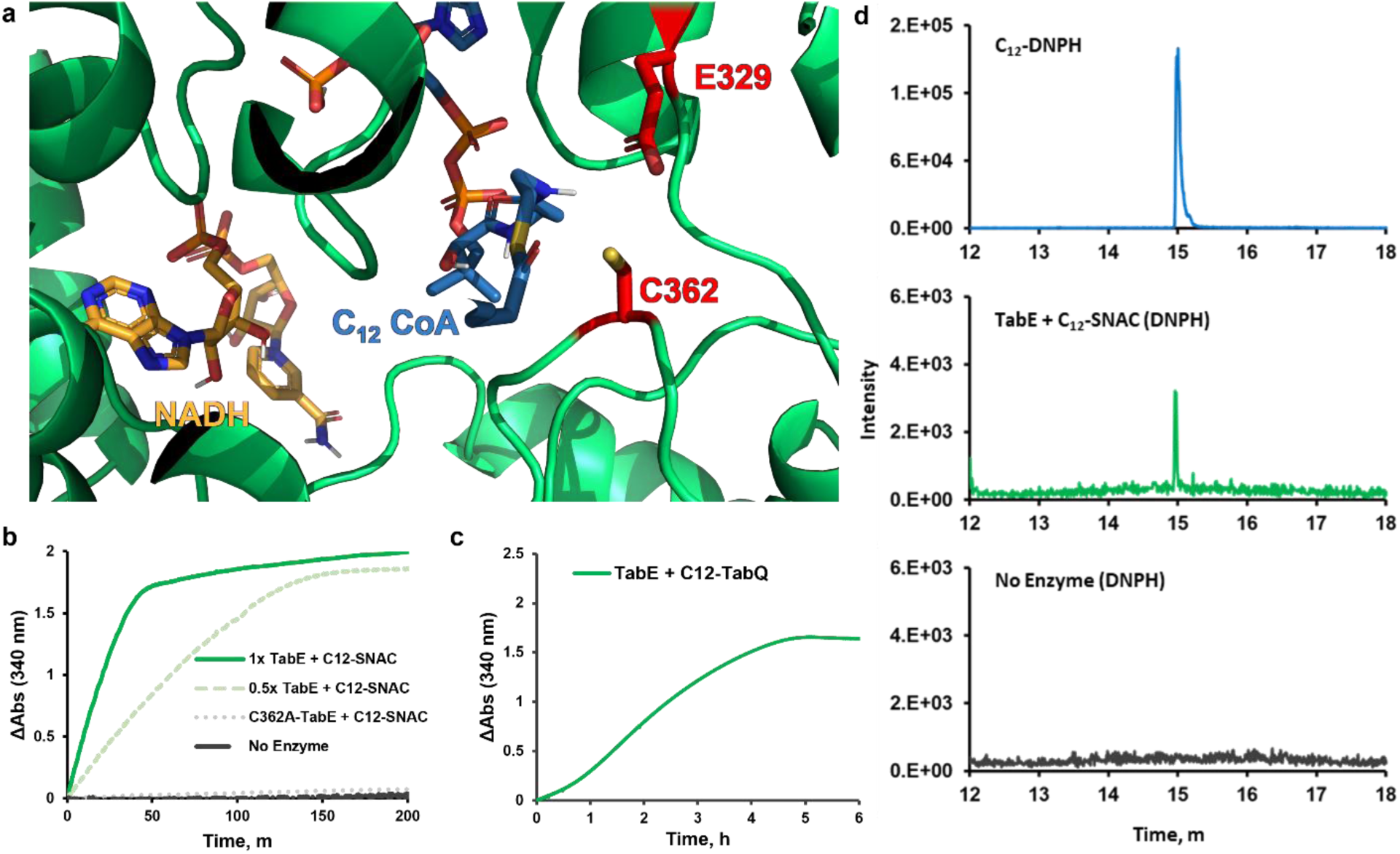
**a)** Chai-1^28^ model of the TabE active site showing the catalytic cysteine (C362) and glutamate (E329) residues (shown in red), NADH (shown in yellow), and C_12_-CoA docked. **b)** Time course for the reaction of TabE (or TabE-C362A) with C_12_-SNAC against a “no enzyme” control. Absorbance was monitored at 340 nm to observe changes in NADH oxidation. **c)** Time course for the reaction of TabE with C_12_-TabQ monitoring NADH oxidation at 340 nm **d)** LC-MS selected ion reporting (363.21 m/z) confirms that C_12_-SNAC is converted to dodecanal by TabE.

At an alignment score of 100 (sequence %ID ≥ 25%), an SSN of the aldehyde/histidinol dehydrogenase homologous superfamily (IPR016161) harbored a putative isofunctional cluster of sequences that included TabE (Figure S4 and Table S1). Every sequence contained in this cluster was annotated as an “aldehyde dehydrogenase” or an “acyl-CoA reductase.” The only sequence in this cluster with any experimental characterization was the NADPH-dependent LuxC^29^ involved in bacterial bioluminescence and fatty alcohol-forming acyl-ACP reductases that likely arose from the fusion of acyl-thioester reductase and aldehyde reductase domains.^30^ This highlights the conserved functional domains and wide functional utility among the “aldehyde dehydrogenase-like” class of enzymes.

Given the revised role of TabJ as an editing thioesterase, one of the remaining enzymes in the Tab pathway must be responsible for offloading the acyl chain from TabQ. Based on the observed clustering of TabE with acyl-CoA reductases in the SSN, this enzyme was putatively assigned as a thioester reductase responsible for this role. As such, TabE was investigated with *in vitro* activity assays using C_12_-SNAC, C_12_-CoA, and C_12_-TabQ as possible substrates and either NADH or NADPH as hydride donors. The activity of TabE was measured by monitoring the rate of NAD(P)H oxidation using the absorbance at 340 nm. When TabE was assayed with NADPH, no significant change in the rate of cofactor oxidation was observed when comparing the reaction with the C_12_-SNAC substrate relative to substrate- and enzyme-free controls (Figure S5 and Figure S6).

Contrastingly, activity patterns observed with NADH were significantly different to those seen for NADPH, suggesting that TabE differentiates between these nicotinamide cofactors. First, the ΔA_340_ observed upon incubating TabE with NADH alone was minimal (Figure S7). When C_12_-SNAC, C_12_-CoA or C_12_-TabQ were included as substrates in this assay, significant increases in the apparent rates of cofactor oxidation were observed (Figure 3b, Figure S7, and Figure 3c, respectively), suggesting that TabE is an NADH-dependent reductive thioesterase. A multiple sequence alignment of TabE with three related ALDH-like proteins (Figure S3) indicated that Cys362 (TabE numbering) was entirely conserved, suggesting a critical role for this residue in the catalytic mechanism of these enzymes. Acyl-CoA reductases often operate via ‘ping-pong’ mechanisms, wherein the growing acyl chain is passed from the CoA thiol to a cysteine in the reductase active-site, and this thioester is reduced in an NAD(P)H-dependent manner.^22^ To evaluate the potential role of C362 as this critical active-site nucleophile, the C362A-TabE mutant was prepared by site-directed mutagenesis, overexpressed, purified, and the activity against the

C_12_-SNAC thioester was compared to the wildtype TabE (Figure 3b). When the C362A-TabE was incubated with this thioester, the observed activity was indistinguishable from controls lacking TabE entirely, demonstrating that C362 is essential for the acyl-ACP thioreduction catalyzed by this enzyme. Based on analogous mechanisms put forth for characterized ALDHs, a reasonable catalytic mechanism for TabE can be proposed (Scheme S3).^31^

Although colorimetric assays unambiguously demonstrated NADH consumption by TabE in the presence of several thioester substrates, these assays offered no direct information about the products of the TabE-catalyzed reaction. To this end, TabE was fed an excess of C_12_-SNAC and NADH, and following overnight incubation, the reaction mixture was treated with 2,4-dinitrophenylhydrazine (DNPH; Brady’s reagent; Scheme S4).^32^ This treatment would convert aldehydes present in this mixture to the corresponding hydrazones, which could be detected using LC-MS. Following incubation and derivatization, the putative C_12_-dinitrophenylhydrazone (Scheme S4) was found to elute with a retention time of approximately 15 minutes, as detected by both absorbance at 360 nm and mass spectroscopy ([M-H]^-^ ion at 363.21 m/z; Figure 3d), consistent with our analysis of an authentic standard of the hydrazone in question and providing additional evidence that TabE catalyzes the conversion of thioesters to aldehydes.

Collectively, the results presented above demonstrate that TabE is a novel stand-alone fatty acyl-ACP reductase that converts acyl-CoAs to aldehydes in an ATP-independent manner. While this is not an entirely unprecedented activity,^22^ TabE seems to exert specificity for NADH, whereas most ATP-independent acyl-ACP reductases preferentially use NADPH as the reductant. LuxC may serve as an important point of comparison: *in vivo*, this enzyme and LuxE (an ATP-dependent acyl-protein synthetase) function together to convert fatty acids into fatty aldehydes. Although the first step catalyzed by the LuxE component of the complex converts the carboxylate substrate into a thioester for subsequent reduction by LuxC, *in vitro* experiments have confirmed that LuxC can also function as a stand-alone reductase to produce aldehydes from acyl-CoAs.^29^ TabE could similarly exist naturally within a hypothetical TabAEJQ multi-enzyme complex where association of each individual polypeptide could facilitate transfer of the product of one enzyme to the active site of the next biosynthetic enzyme.

Importantly, TabE does not have a functional analog in the Tam pathway. While TamH does harbor a thioreductase domain, TabE shares only 15.5 % sequence identity with this protein. No aldehyde dehydrogenase or acyl-CoA reductase domain was detected in TamH using the InterProScan tool. While there is alignment of the catalytic cysteines and oxyanion-stabilizing asparagines between TabE and the thioreductase domain of TamH, the latter enzyme appears to utilize arginine in place of glutamate as a general base (Figure S3). Although the transformations catalyzed by the reductase domains of TamH and TabE are similar, TabE appears to have evolved as a discrete acyl-ACP reductase toward the biosynthesis of **6** in *Streptomyces*. The observation that two seemingly unrelated enzymes have evolved to carry out the same biosynthetic role could agree with previous suggestions that the machinery involved in bipyrrolic NP production emerged, at least in part, as a result of convergent evolution.^6^

### TabA is a ω-transaminase that catalyzes the final step in biosynthesis of **6**

In the context of **3** biosynthesis, the product of the TabE-catalyzed reaction, dodecanal, is hypothetically converted to **6** by TabA – a putative class III fold ω-transaminase (NCBI accession: WP_078565688.1). These enzymes form covalent adducts between a conserved lysine residue within the active site and a PLP cofactor (Figure 4a). The holoenzyme then uses an amine donor (typically an amino acid) to catalyze reductive amination of aldehydes or ketones.^34^

**Figure 4.**
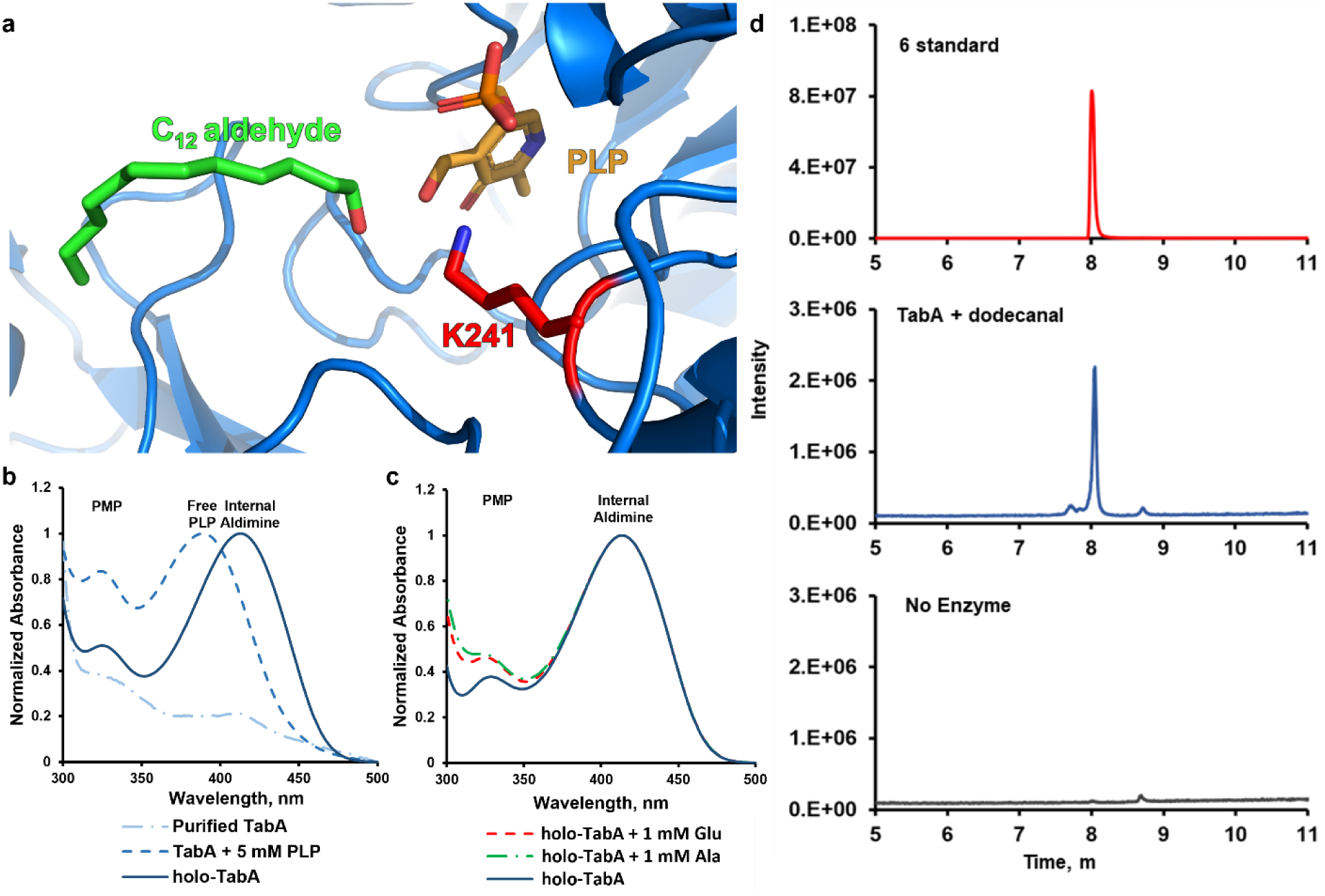
**a)** Chai-1^28^ model of the TabA active site showing the lysine residue which forms a covalent adduct with PLP (shown in yellow) docked non-covalently within the active site. **b**) Spectrophotometric analysis of the binding and release of PLP, PMP and formation of internal aldimine. **c)** Spectrophotometric analysis of TabA amine donor showing release of PMP upon addition of L-glutamate and L-alanine. **d)** LC-MS selected ion reporting (186.36 m/z) chromatograms for the conversion of dodecanal to DDA by TabA against a “no enzyme” control.

The ability of TabA to bind PLP and amino acid amine donors was assessed spectrophotometrically. Incubating TabA with PLP and subsequently removing excess cofactor resulted in an increased absorbance at 415 nm relative to the *apo*-TabA, indicating formation of the PLP-TabA aldimine (*holo*-TabA) (Figure 4b).^8^ Additionally, incubating *holo*-TabA with L-glutamate or L-alanine resulted in the release of pyridoxamine (PMP), as indicated by an increase in the normalized absorbance at 320 nm.^8^ This indicated that both L-glutamate and L-alanine can be used by TabA as amine donors (Figure 4c). To confirm that the observed absorbance changes corresponded to enzyme-catalyzed transamination, dodecanal and alanine were incubated with

TabA for 48 hrs before the reaction was quenched and analyzed LC-MS. The [M+H]^+^ ion (186.36 m/z) corresponding to the fatty amine product **6** was detected at an elution time of 8 minutes and further confirmed using an authentic standard of **6** (Figure 4d).

With the function of TabA confirmed, the *tam* BGC was evaluated for similar functionalities. TabA shares 23.2% sequence identity with the transaminase domain of TamH. As described above, TamH contains functional elements of both TabE and TabA in a single polypeptide chain, but the thioreductase and aminotransferase domains of this protein have low similarities to the individual enzymes that perform these functions in the Tab pathway, suggesting the Actinobacteria employ biosynthetic logic that is distinct from Proteobacteria in the biosynthesis of tambjamine tails.

### Enzymes from outside the tab BGC are likely required for BE-18591 tail biosynthesis

The enzymes present in the Tam pathway are specialized for recruiting and modifying a fatty acid from primary metabolism. Contrastingly, the Tab pathway is a putative hybrid primary/secondary metabolism pathway that constructs a dedicated on-pathway intermediate using enzymes involved in fatty acid biosynthesis.^6^ Bailey and West recently reviewed the “crosstalk” between primary fatty acid metabolism and polyketide biosynthetic pathways, lending insight into the co-evolution of primary and secondary metabolism.^35^ Reynolds and co-workers found that the *Streptomyces* fatty acid biosynthetic acyltransferase, FabD, does not discriminate between ACPs but the ketosynthase, FabH, does.^36^ This would necessitate the presence of dedicated ketosynthases such as TabP or RedP (undecylprodigiosin biosynthesis) in their respective pathways. In a RedP deletion mutant of *Streptomyces coelicolor*, there was an 80% reduction in total prodiginine production, and production levels were entirely restored by complementation with FabH from *Streptomyces glaucenscens*.^37^ Subsequent work from Reynolds demonstrated that ACP variants of RedQ (an orthologue of TabQ) can be processed by the fatty acid biosynthetic ketoreductase (FabG), dehydratase (FabA), and enoylreductase (FabI) in *S. coelicolor*.^38,39^ High similarities between the fatty acid biosynthetic enzymes in *S. coelicolor* A3(2) and *S. albus* NRRL B-2362 and between the Tab and Red enzymes (Tables S4 – S5) suggests the prodiginine-fatty acid crosstalk that is operative in undecylprogiosin biosynthesis might also be operative in the biosynthesis of **3**. As such, an updated biosynthetic scheme for **6** biosynthesis has been constructed based on the findings presented here and our hypotheses regarding the dual role of enzymes from primary fatty acid biosynthesis (Scheme 2).

## CONCLUDING REMARKS

Here, four enzymes from the Tab biosynthetic pathway have been characterized to better understand the biosynthetic logic that different phyla employ for tambjamine production. TabQ acts as an acyl-carrier protein to anchor a growing acyl chain. TabJ, a type II editing thioesterase, aids in selecting for a C_12_ acyl chain by cleaving aberrant acyl chains from TabQ. The resulting C_12_ acyl chain is reductively offloaded from TabQ by the acyl-ACP reductase, TabE, to afford the free C_12_ aldehyde. The aldehyde is then converted into the C_12_ amine by the ω-transaminase, TabA, which will become the tail fragment of tambjamine BE-18591.

Our findings revealed novel biosynthetic logic toward the biosynthesis of tambjamines in Actinobacteria. *In vitro* characterization of the novel enzymes in the Tab pathway implicated in the biosynthesis of **3** have suggested the need for a revised biosynthetic scheme for this tambjamine. While the chemistries employed to process the fatty acid in both **2** and **3** biosynthetic pathways are similar, the evolutionary origins of these pathways are clearly disparate. The findings herein thus suggest the independent evolution of long-tailed tambjamines in Actinobacteria. Our future efforts will focus on tracing the evolutionary lineage of the fragments of tambjamine and prodiginine gene clusters to suggest mechanisms of evolution (eg. horizontal gene transfer) and identify the selective pressures that gave rise to the production of bipyrrolic alkaloids in the producing organisms. Several studies have honed in on the ability of tambjamines to transport chloride across lipid bilayers highlighting this as one of the primary mechanisms of bioactivity and therapeutically relevant.^40,41,42,43,44^ Probing the structural features of tambjamines that contribute to biological activity will put into context why the producing organisms converge on common structural motifs. Collectively with this work, these investigations will unravel how Nature selects for and constructs potently bioactive molecules.

## AUTHOR INFORMATION

### Author Contributions

NLG designed experiments, performed computational analyses, wet lab experiments, and contributed to drafting and editing. YF and EHP helped perform wet lab experiments. ACR and GWH contributed to experimental design, editing, and provided funding for the work herein. All authors have given approval to the final version of the manuscript.

## ACKNOWLEDGMENT

NLG is the grateful recipient of a Natural Sciences and Engineering Research Council (NSERC) postgraduate scholarship. We also gratefully acknowledge the labs of D. Zechel (Queen’s University) and Z. Jia (Queen’s University) for support and resources on this project and D. Campopiano (University of Edinburgh) for productive discussions on characterizing biosynthetic enzymes.

## METHODS

### General experimental protocols

All NMR spectra were recorded on Bruker Avance 300 MHz or Bruker Avance NEO 500 MHz spectrometers in CDCl_3_. High resolution mass spectra were recorded on a Thermo Scientific Orbitrap Velos Pro Easy-nLC/HESI Hybrid Ion Trap-Orbitrap mass spectrometer by direct injection, a cone voltage of 175 V, and scanning 40 – 520 m/z in positive ion mode. and LC-MS data were collected on a Waters H-Class UPLC-MS with a C_18_ column (Waters CORTECS UPLC T3; 2.1 mm x 100 mm; 1.6 μm particle size; 120 Å pore size). UV-active analytes were detected by photodiode array absorbance from 200 – 600 nm. Electrospray ionization into a single quadrupole mass spectrometer at a 0.5 kV electrospray voltage scanning 100 – 2048 m/z. UV-Vis data were collected on a Cary 3500 UV-Vis spectrophotometer (Agilent Technologies). Microplate data were collected on a Synergy H1 microplate reader (BioTek) using UV-Star 96-well plates (Greiner Bio-One). Expression constructs were synthesized commercially (Biomatik) into the pET-28a expression system and transformed into chemically competent *Escherichia coli* BL21 (DE3). The gene for TabJ was re-cloned into a modified pET-16b vector (HT29) with a maltose-binding protein fusion tag to improve solubility (Zongchao Jia, Queen’s University).^45^

### Cloning of TabJ-MBP fusion expression construct

Target sequence was amplified by Phusion PCR (New England Biolabs) using the primers listed in Table S2. The amplified sequences were ligated into HT29 vector (derived from pET16b) with ampicillin resistance marker (linearized with NdeI and EcoRI) using the iVEC3 *in vivo* cloning system (*E. coli* K-12 MG1655 ΔhsdRΔendAΔrecA) following the previously reported protocol.^46^ The resulting construct was extracted using a Monarch Plasmid Mini Prep kit (New England Biolabs). Successful cloning was confirmed by DNA sequencing (Plasmidsaurus). The resulting plasmid DNA was transformed into chemically competent *E. coli* DH10β for storage and into *E. coli* BL21 (DE3) for over-expression.

### Protein overexpression and purification

Overnight cultures of *E. coli* BL21 (DE3) transformed with the appropriate construct in 10 mL LB broth (Bioshop) containing 50 μg/mL of the appropriate selection antibiotic was inoculated and incubated at 37 °C with shaking at 180 rpm. On the following day, 1 L of LB with antibiotic was inoculated with the overnight culture and incubated at the same condition until the OD_600_ reached 0.4-0.8. Over-expression was then induced by adding 0.5 M isopropyl ß-D-1-thiogalactopyranoside (IPTG) and incubating at 16 °C with shaking at 180 rpm for 18-24 hrs.

The cells were pelleted by centrifugation and resuspended in 40 mL of lysis buffer (20 mM Tris pH 7.6, 100 mM NaCl, 10% v/v glycerol). Cells were lysed under high pressure using an Emulsiflex-C5 (Avestin). Lysate was centrifuged again to remove debris and the supernatant was loaded on 5 mL of nickel sepharose resin (Cytiva). Resin was washed twice with 10 column volumes of wash buffer (20 mM Tris pH 7.6, 100 mM NaCl, 20 mM imidazole, 10% v/v glycerol). Protein of interest was eluted with 5 column volumes of elution buffer (20 mM Tris pH 7.6, 100 mM NaCl, 500 mM imidazole, 10% v/v glycerol).

Eluent was concentrated using 15 mL Amicon centrifugal filters (Millipore) with the appropriate molecular weight cutoff. Sample was exchanged into storage buffer (40 mM Tris pH 7.6, 100 mM NaCl, 10% glycerol) by size exclusion with PD10 columns (Cytiva). Samples were quantified using a nanophotometer (Implen NP80), and flash frozen in liquid nitrogen for storage at −80 °C.

### TabQ loading reaction

Reaction mixtures contained 50 μM apo-TabQ, 150 μM acyl-CoA, and 2 μM Sfp. Each reaction was brought to a final volume of 100 μL in 50 mM Tris buffer pH 7.5, 500 μM TCEP, and 2.5 mM MgCl_2_ and was incubated for 2 hrs at 37 °C with shaking. Excess acyl-CoA was removed using an Amicon centrifugal filter (Millipore) with a 3 KDa molecular weight cutoff. Loading was confirmed by LC-MS (gradient 8.5:0.5:1 H_2_O:acetonitrile:1% formic acid to 9:1 acetonitrile:1% formic acid over 25 mins)

### Synthesis of acyl S-NAC thioesters

A solution of corresponding fatty acid (Sigma) was prepared by dissolving 10 mmol in 30 mL dichloromethane (Fischer) and cooled to 0 °C for ten minutes. 1-ethyl-3-(3-dimethylaminopropyl) carbodiimide (EDCI, 10 mmol), 4-dimethylaminopyridine (DMAP, 10 mmol) and *N*-acetylcysteamine (10 mmol) were added and the mixture was left stirring overnight at room temperature. The reaction mixture was concentrated under reduced pressure and extracted with saturated aqueous ammonium chloride and ethyl acetate. Organic fractions were pooled, dried over anhydrous magnesium sulfate, and further purified by flash chromatography (Büchi Pure C-810 Flash) using silica columns (Büchi FlashPure EcoFlex Silica 50 μm 120 g) with 1:1 ethyl acetate:hexanes to provide the desired SNAC thioester derivative. The identity of the compounds was confirmed by LC-MS (gradient 8.5:0.5:1 H_2_O:acetonitrile:1% formic acid to 9:1 acetonitrile:1% formic acid over 30 mins) and NMR spectroscopy (appendix A in the supplementary information).

### Ellman’s assay of TabJ activity

TabJ hydrolytic activity assay performed in 96-well plates by monitoring absorbance at 412 nm on a microplate reader at 30 °C with orbital shaking at 180 cpm. Measurements were recorded every 2 mins for the first hour then every hour for 23 hrs. Reaction mixtures contained 1μM TabJ, 100 μM DTNB, and 500 μM substrate (from DMSO stock for acyl-SNACs and Milli-Q H_2_O stock for acyl-CoAs) or 10 μL of the TabQ loading reaction. Each well was brought to a final volume of 200 μL with TabJ reaction buffer (100 mM Tris pH 7.4, 20 mM NaCl). The absorbance of the TNB anion begins to quench over the course of 24 hours under acyl-SNAC, - CoA, and -TabQ experimental conditions. To account for this, the absorbance values presented for the C_12_-TabQ Ellman’s assay with TabJ were processed by adding the slope of the “No Enzyme” negative control to both test and control conditions. The raw data for the plot in Fig. 3e has been provided in the supplementary information appendix C.

### NAD(P)H oxidation assay of TabE activity

Reaction mixtures of 500 μM NAD(P)H, 5 μM TabE, and 30 μL TabQ loading reaction or 500 μM C_12_-SNAC/CoA were brought to a final volume of 1 mL in TabE reaction buffer (50 mM Tris pH 7.5, 100 mM NaCl, 10 mM MgCl_2_, 1 mM DTT) and were analyzed by UV-Vis spectrophotometry. The reaction was monitored at 340 nm and 30 °C overnight.

### TabE reaction product derivatization

NADH oxidation assay with C_12_-SNAC was scaled up to 2 mL and incubated overnight at 30 °C. To the reaction mixture, 2 mL MeOH, 1 mL 5M HCl and 100 mg DNPH were added, and the mixture was heated to 60 °C for 3 hrs. The reaction mixture was filtered and 1 μL injected onto the UPLC-MS and separated using a gradient of 57:40:3 H_2_O:acetonitrile:1% formic acid to 32:65:3 H_2_O:acetonitrile:1% formic acid over 9 mins to 97:3 acetonitrile:1% formic acid over 4 mins and finally isocratic 97:3 acetonitrile:1% formic acid for 7 mins. The DNPH derivative was detected by monitoring the [M-H]^-^ peak at 363.21 m/z and absorbance at 360 nm.

### TabA amine donor uptake

An aliquot of TabA in storage buffer (∼4 mg/mL) with 5 mM PLP added was incubated at 30 °C overnight with orbital shaking at 180 rpm. The following day, excess PLP was removed using a PD-10 desalting column (Cytiva) to afford *holo*-TabA. Separate aliquots of *holo*-TabA were incubated with 1 mM L-glutamate, L-alanine or no additive (control) at 30 °C overnight with orbital shaking at 180 rpm. Before and after each treatment, the absorbance of each aliquot was recorded from 300 - 500 nm.

### Assay of TabA activity

A 5 mL reaction mixture of 50 nM TabA, 100 μM PLP, 2.5 mM dodecanal and alanine in TabA reaction buffer (50 mM HEPES, pH 8.0) was incubated at 30 °C for 48 hours. The reaction was quenched with 1:1 acetonitrile:0.01% trifluoroacetic acid, filtered and 1 μL injected onto the LC-MS using a gradient of 85:5:10 H_2_O:acetonitrile:1% formic acid to 9:1 acetonitrile:1% formic acid over 13 mins. The DDA product was detected by monitoring at [M+H]^+^ m/z 186.35.

### Bioinformatics and in silico methods

Gene cluster comparison and visualization was performed by inputting GenBank files of each BGC to the clinker tool in CAGECAT^47^. Output was manually annotated and trimmed.

All protein structure visualization was performed using PyMOL 2.4 (https://pymol.org/2). Root-mean-square deviation values were calculated using the Alignment plugin in PyMOL2. Substrate tunnels were computed using the Caver 3.0.3 plugin for PyMOL2.^48^ Substrate/co-factor ridged docking was performed using AutoDockTools4^49^.

The amino acid sequence of TabE from *S. albus* NRRL B-2362 was analyzed with the InterProScan tool to identify the InterPro family of any functional domains present. TabE was annotated as a member of the aldehyde/histidinol dehydrogenase homologous superfamily (IPR016161). The EFI-EST was used to perform an all-by-all BLAST for the sequences found within the UniRef50 database that are members of each of this protein family. All sequence similarity networks (SSNs) were visualized in Cytoscape 3.10.1.^50^

Pairwise sequence alignments were performed using EBMOSS Needle and Multiple sequence alignments were generated using Clustal Omega with default settings. Alignments were visualized in MView.^51^ Alignments querying *tam* genes were performed on the orthologues from *Pseudoalteromonas citrea*.

### Site-directed mutagenesis of TabE

The C362E-TabE mutant was generated using a Q5 site-directed mutagenesis kit (New England Biolabs). PCR was performed using the primers listed in Table S3 and pET28a-TabE as the template. Amplification was confirmed by 1% agarose gel electrophoresis. Plasmid mutation was confirmed by sequencing (Plasmidsaurus) and transformed into chemically competent *E. coli* BL21 (DE3) for heterologous over-expression and purification.

---

**Scheme 1.**
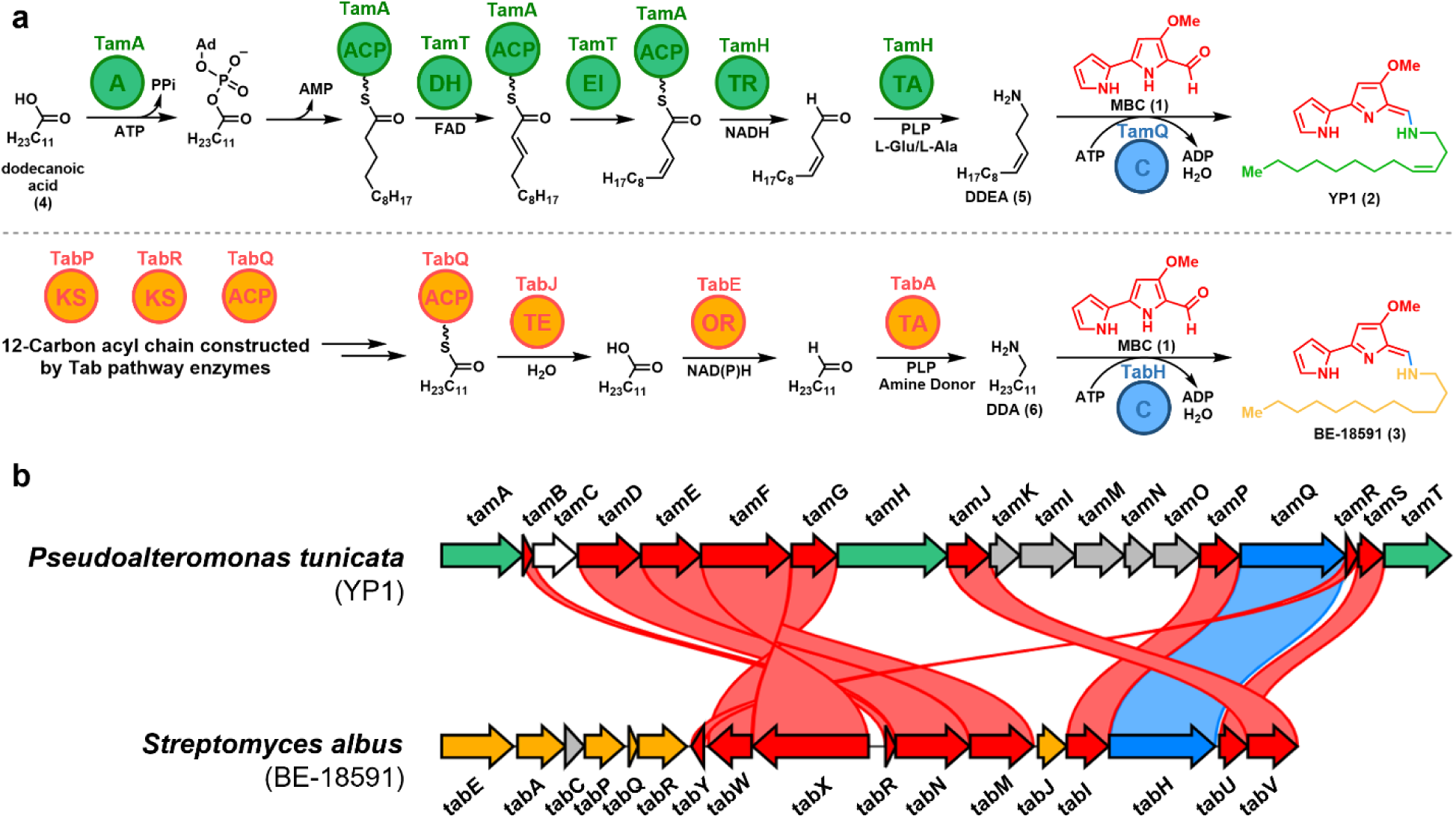
Comparison of YP1 (**1**) and BE-18591 (**2**) biosynthesis. **a)** Proposed biosynthetic schemes for the construction of the tails of YP1 (**2**) (top) and BE-18591 (**3**) (bottom). Protein functions are as follows: A, Adenylation; ACP, acyl carrier protein; C, condensation; DH, dehydrogenase; EI, enoyl isomerase; KS, ketosynthase; OR, oxidoreductase; TA, transaminase; TE, thioesterase; TR, thioreductase. **b)** Comparative analysis of the *tam* and *tab* BGCs. Genes with >30% BLAST similarity are connected. Gene colors match the fragment of the NP that they encode: red genes for MBC, green genes for DDEA (**5**), orange genes for DDA (**6**), and blue genes for condensation.

**Scheme 2.**
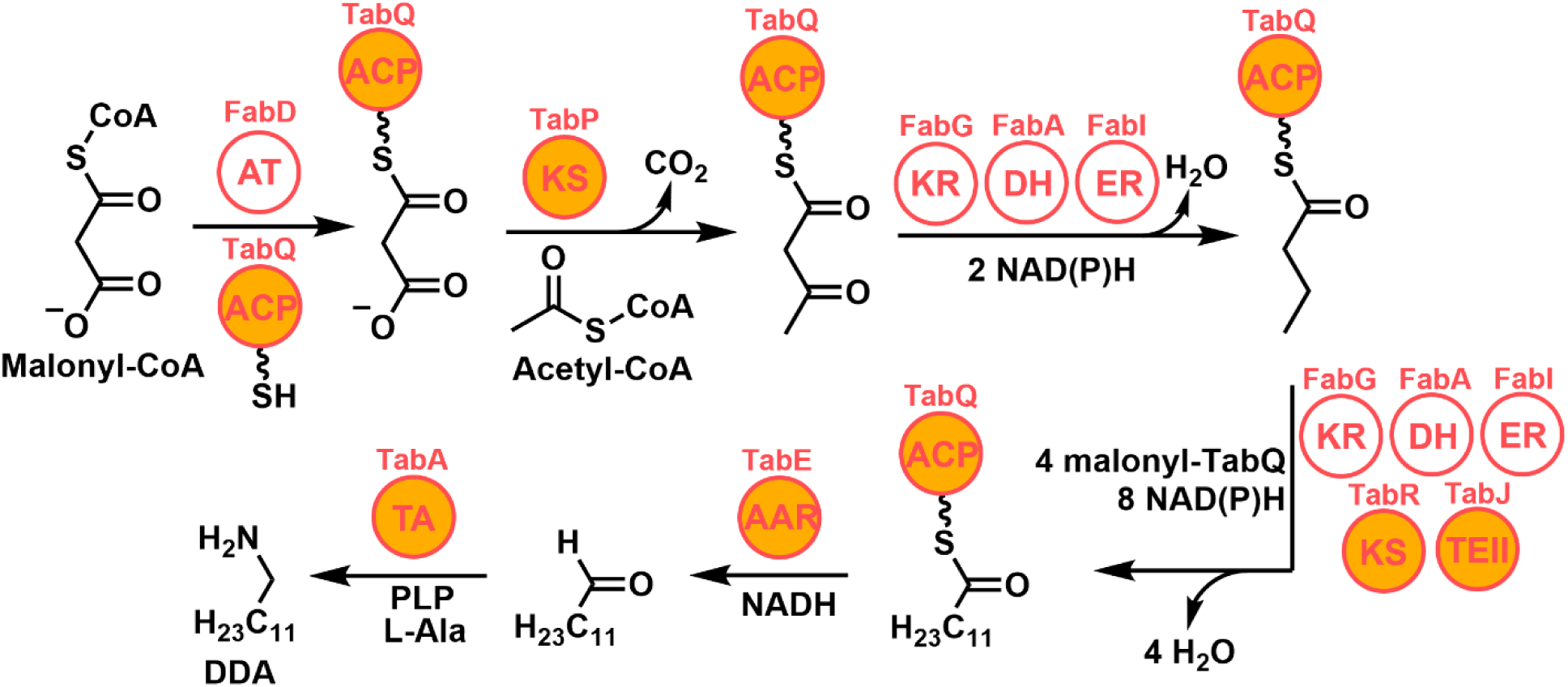
A refined scheme for the biosynthesis of DDA – the precursor to the tail of BE-18591 (2) – by the Tab pathway. AAR, acyl-ACP reductase; AT, acyl transferase; ACP, acyl carrier protein; DH, dehydratase; ER, enoyl reductase; KR, ketoreductase; KS, ketosynthase; TA, transaminase; TEII, type II thioesterase.

