## Supplemental Information for "Characterization of a novel fatty acid-modifying pathway toward the biosynthesis of tambjamine BE-18591 in *Streptomyces*"

### Table of Contents

**Figure S1: Multiple sequence alignment of TabQ** .......... S3

**Figure S2: Multiple sequence alignment of TabJ** .......... S3

**Scheme S1: Chemical synthesis of acyl S-NAC thioesters** .......... S4

**Scheme S2: Hydrolysis of acyl S-NAC thioester by TabJ and transthiolation with Ellman's reagent** .......... S4

**Figure S4: Multiple sequence alignment of TabE** .......... S4

**Figure S5: Sequence similarity network of the aldehyde/histidinol dehydrogenase superfamily (IPR016161)** .......... S5

**Table S1: Putative functions of enzymes within clusters from Fig. 4c** .......... S5

**Figure S5: NADPH kinetic time-course for TabE in the absence of substrate.** .......... S6

**Figure S6: NADPH kinetic time-course for TabE with $C_{12}$-SNAC** .......... S7

**Figure S7: Kinetic time-course of TabE with $C_2$-CoA and $C_{12}$-CoA against a no-substrate control** .......... S7

**Scheme S3: Putative mechanism of TabE** .......... S8

**Scheme S4: Derivatization of dodecanal for LC-MS analysis** .......... S8

**Table S2: Primers for cloning of TabJ-MBP construct** .......... S8

**Table S3: Primers for site-directed mutagenesis of TabE to generate C362A mutant** .......... S8

**Table S4: Pairwise analysis of primary and secondary fatty acid metabolic enzymes genes in *S. albus* NRRL B-2362 *and S. coelicolor* A2(3)** .......... S9

**Table S4: Pairwise analysis of primary fatty acid biosynthetic enzymes against their secondary metabolism counterparts** .......... S9

**References** .......... S9

**APPENDIX A: Acyl-SNAC NMR, HR-ESI-MS, UPLC-PDA data** .......... S10

**APPENDIX B: Plasmid maps and sequencing** .......... S21

**APPENDIX C: TabJ $C_{12}$-TabQ assay raw data** .......... S27

**APPENDIX D: Protein purification SDS-PAGE** .......... S29

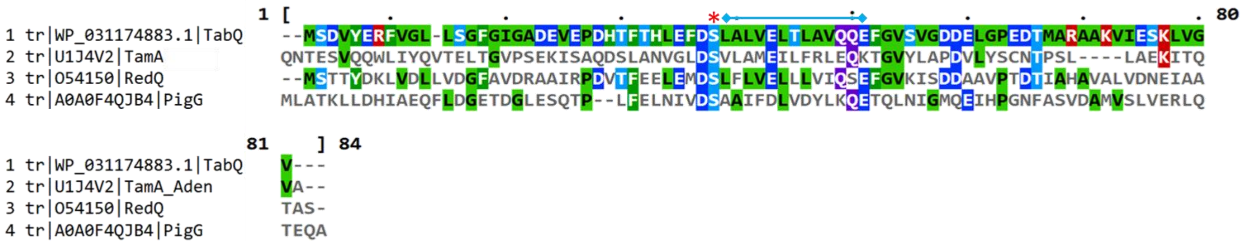


**Figure S1.** Multiple sequence alignment of TabQ with the acyl-carrier domain of TamA, an orthologous protein from the undecylprodigiosin biosynthetic pathway, RedQ, and an acyl carrier protein from the prodigiosin biosynthetic pathway, PigG. The conserved serine residue is denoted by a red asterisk and the putative recognition helix by a blue line capped with diamonds. Alignment was performed using Clustal Omega[1] and visualized in MView.

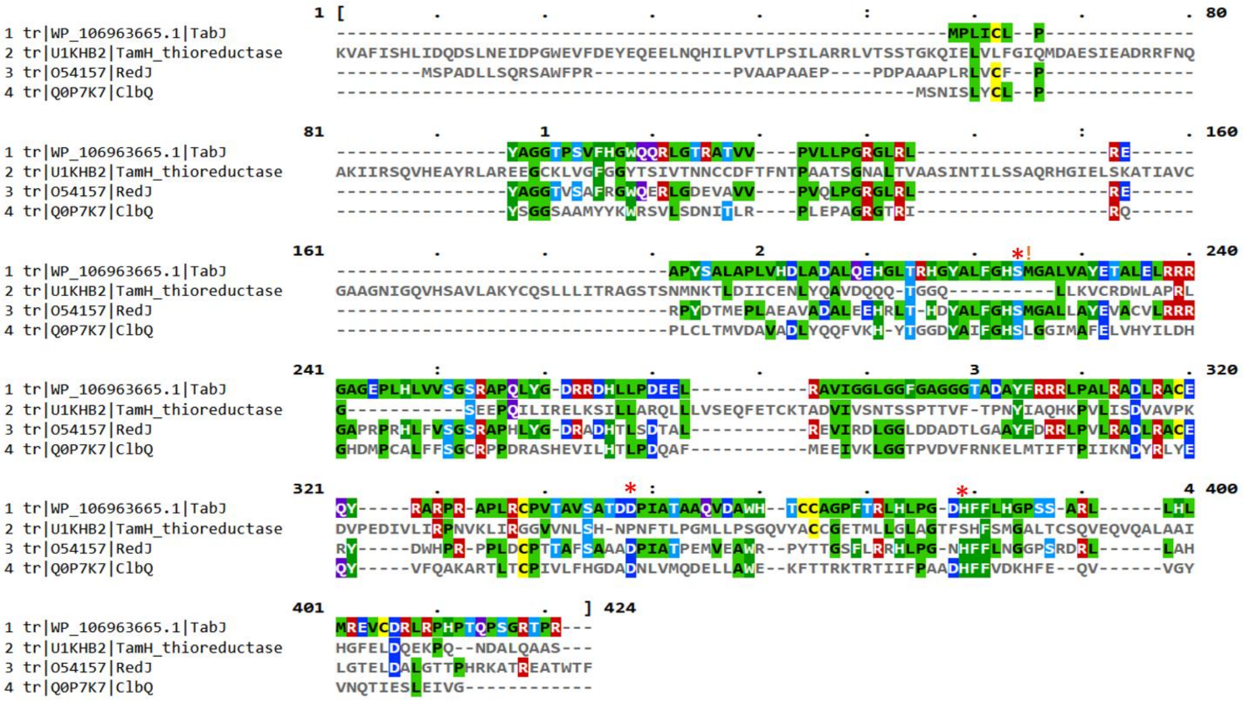


**Figure S2.** Multiple sequence alignment of TabJ with the thioreductase domain of TamH, an orthologous type II thioesterase from the undecylprodigiosin biosynthetic pathway, RedJ, and a well-characterized type II thioesterase from the coelibactin biosynthetic pathway, ClbQ. The conserved residues of the Ser-His-Asp catalytic triad are denoted by red asterisks. Alignment was performed using Clustal Omega and visualized in MView.

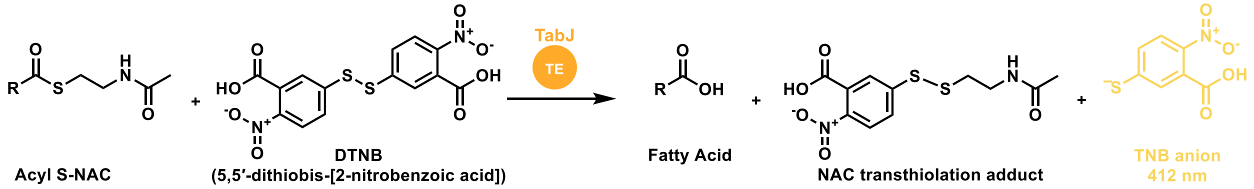


**Scheme S1.** Hydrolysis of acyl S-NAC thioester by TabJ and transthiolation with Ellman's reagent. DTNB undergoes a transthiolation with free thiols releasing a TNB anion that is bright yellow and absorbs strongly at 412 nm.

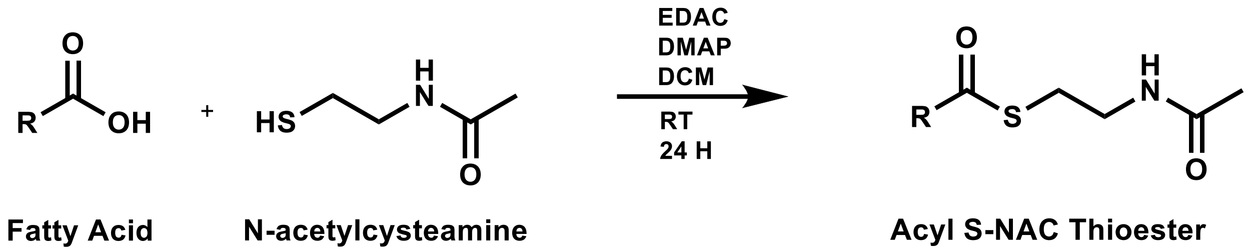


**Scheme S2.** Chemical synthesis of acyl S-NAC thioesters. EDAC, 1-ethyl-3-(3-dimethylaminopropyl)carbodiimide; DMAP, 4-dimethylaminopyridine; DCM, dichloromethane. Adapted from Guntaka et al. 2017[2].

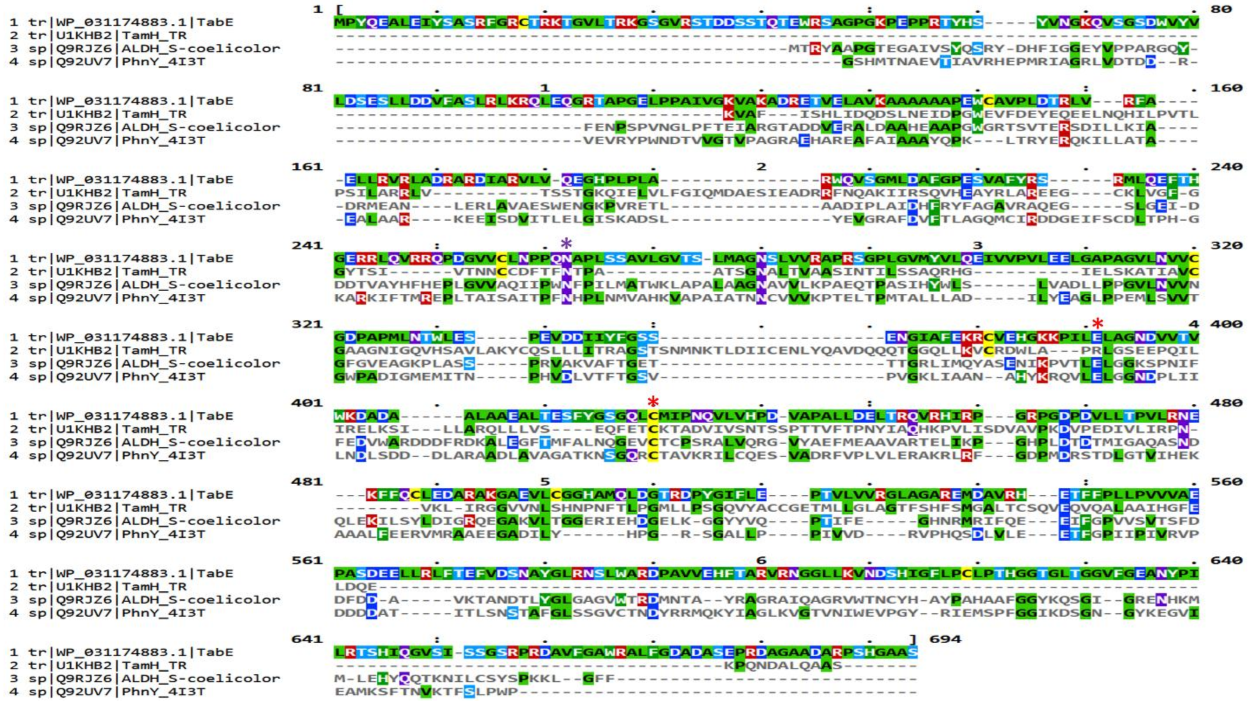


**Figure S3.** Multiple sequence alignment of TabE with the thioreductase domain of TamH, a characterized aldehyde dehydrogenase from *S. coelicolor*, and PhnY, an aldehyde dehydrogenase from *Sinorhizobium meliloti*. The conserved catalytic residues denoted by red asterisks and the oxyanion stabilizing asparagine residue is denoted by a purple asterisk. Alignment was performed using Clustal Omega and visualized in MView.

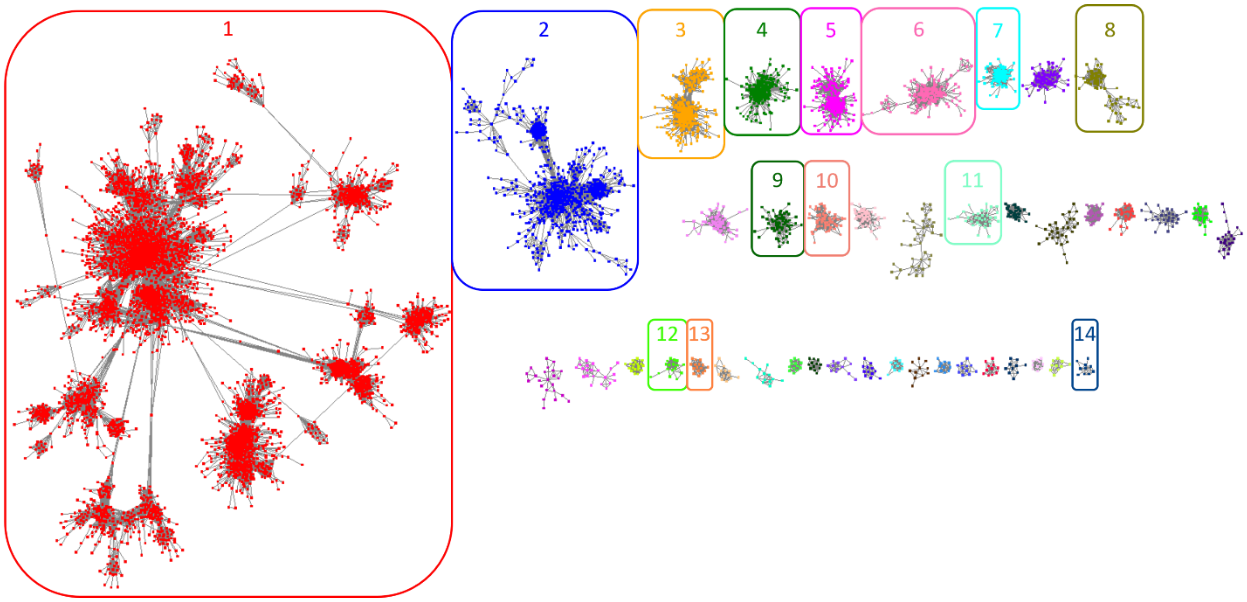


**Figure S4.** Sequence similarity network of the aldehyde/histidinol dehydrogenase superfamily (IPR016161). Each node represents an enzyme sequence or multiple enzyme sequences with greater than or equal to 50% sequence identity across at least 80% of the sequence. An edge (line) drawn between any two nodes indicated at least 25% ID between the sequences in each node and corresponds to an alignment score of 100. A putative isofunctional cluster of TabE-like enzymes is located in cluster 12. Putative representative annotations of each cluster can be found in Table S1.

**Table S1.** Putative functions of enzymes within clusters from Figure S5 based on representative SwissProt annotations of enzymes within each highlighted cluster. Putative TabE sequence is found in cluster 12.

| Cluster | Representative SwissProt Annotation |
|---|---|
| 1 | Aldehyde Dehydrogenase |
| 2 | Histidinol dehydrogenase |
| 3 | Gamma-glutamyl phosphate reductase |
| 4 | Succinate semialdehyde dehydrogenase |
| 5 | Alpha-ketoglutaric semialdehyde dehydrogenase |
| 6 | alcohol-aldehyde dehydrogenase |
| 7 | Delta-1-pyrroline-5-carboxylate dehydrogenase |
| 8 | N-succinylglutamate 5-semialdehyde dehydrogenase |
| 9 | Acetaldehyde dehydrogenase (acetylating) |
| 10 | Aromatic carboxylate degrading enzyme |
| 11 | Sulfoacetaldehyde dehydrogenase |
| **12** | **Putative TabE-like enzymes** |
| 13 | Succinate-semialdehyde dehydrogenase (acetylating) |
| 14 | Long-chain acyl-protein thioester reductase |

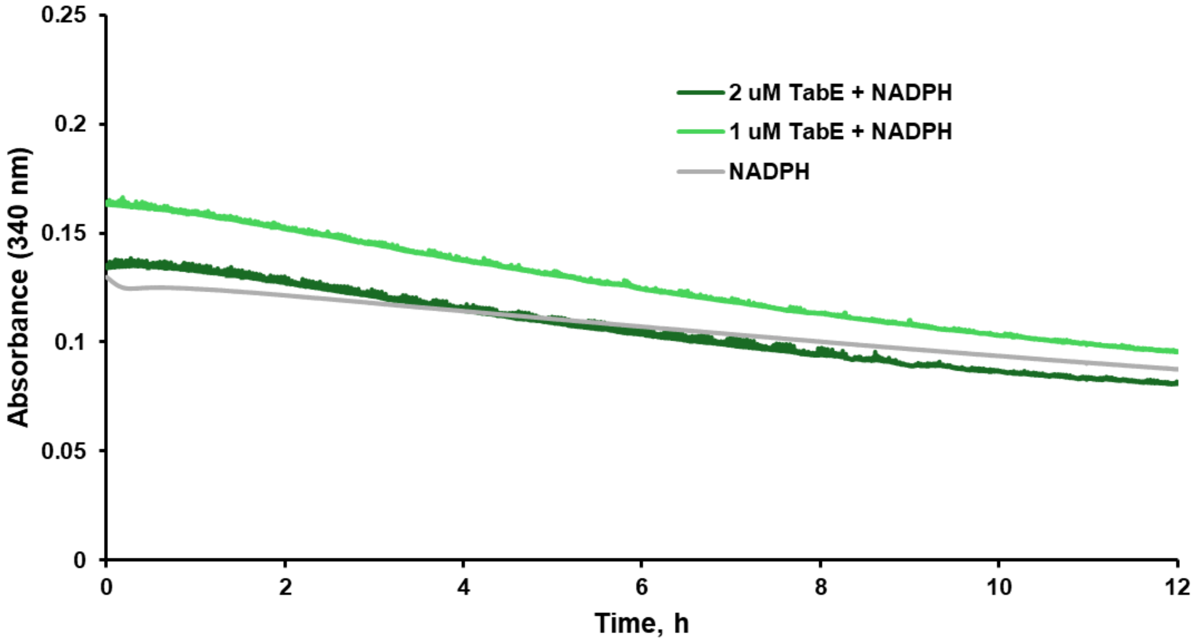


**Figure S5.** NADPH kinetic time-course for TabE in the absence of substrate. Absorbance is monitored at 340 nm to observe potential differences in the rates of NADPH oxidation indicated by a decrease in the absorbance.

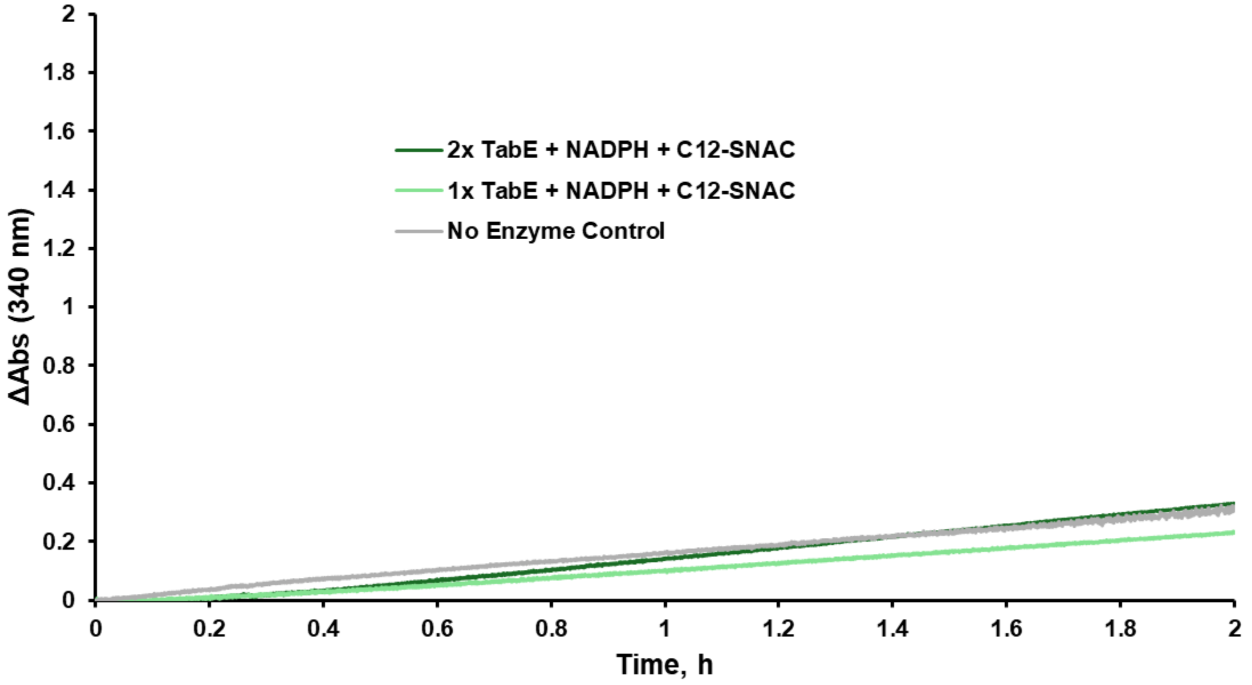


**Figure S6.** NADPH kinetic time-course for TabE with $C_{12}$-SNAC. Absorbance is monitored at 340 nm to observe potential differences in the rates of NADPH oxidation indicated by a decrease in the absorbance.

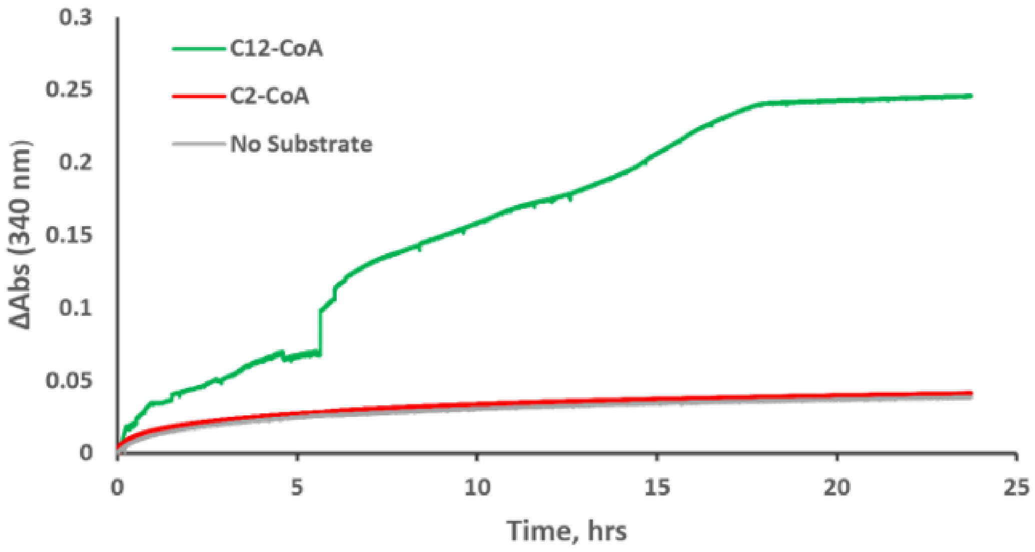


**Figure S7.** Kinetic time-course of TabE with $C_2$-CoA and $C_{12}$-CoA against a no-substrate control. Absorbance is monitored at 340 nm to observe differences in NADH oxidation.

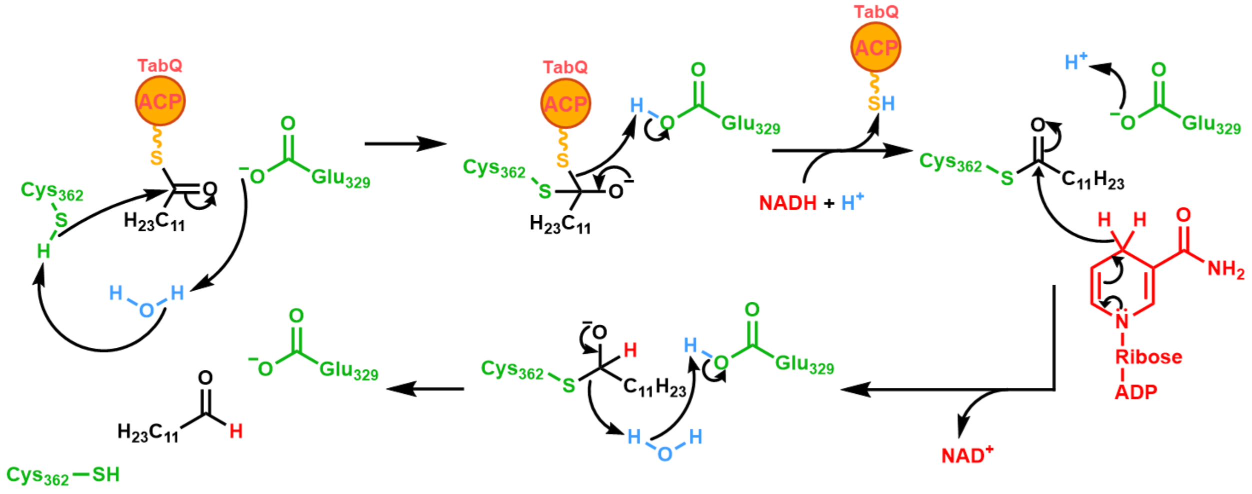


**Scheme S3.** Putative mechanism of TabE. C362 is acts as a crucial nucleophile in the reduction of the acyl-TabQ through a thioacyl-enzyme intermediate.

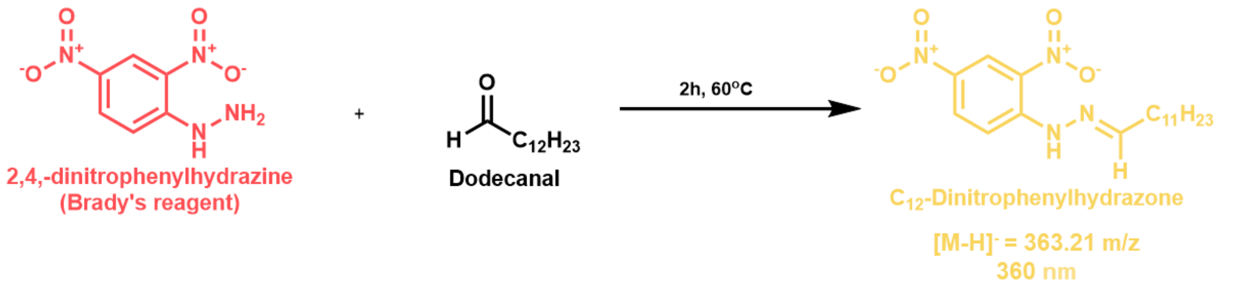


**Scheme S4.** Derivatization of dodecanal for LC-MS analysis. Brady's reagent provides a stable ionizable moiety for aldehydes to be analyzed using negative ion mass spectrometry.

**Table S2.** Primers for cloning of TabJ-MBP construct. Binding regions to the target sequence are in bold and underlined.

| Clone | Primer | Sequence (5'-3') |
|---|---|---|
| HT29-TabJ | tabJ-MBP-F | ACCTGTATTTTCAGTCTGGAGGATCCCATA**ATCATCATCATCATCACAGCA** |
| | tabJ-MBP-R | GTCAGTCACGATGCGGCCGCTCGAGGAATT**GATCTCAGTGGTGGTGGTG** |

**Table S3.** Primers for site-directed mutagenesis of TabE to generate C362A mutant. Mutating nucleotides are in bold and underlined.

| Mutation | Primer | Sequence (5'-3') |
|---|---|---|
| C362A | TabE_C362A_F | TGGACAATTA**GCG**ATGATCCCCAATCAGGTTTTAG |
| | TabE_C362A_R | GACCCATAGAAGCTCTCG |

**Table S4.** Pairwise analysis of primary and secondary fatty acid metabolic enzymes genes in *S. albus* NRRL B-2362 *and S. coelicolor* A2(3).

| | Primary Fatty Acid Biosynthesis | | | | Secondary Metabolite Biosynthesis | | | |
|---|---|---|---|---|---|---|---|---|
| Function | Orthologue Name | *S. albus* | *S. coelicolor* | BLAST %ID | Orthologue Name | Tab Enzyme | Red Enzyme | BLAST %ID |
| Acyl-carrier protein | FabC | WP_016468813.1 | WP_003976417.1 | 87.8 | Q | WP_016467649.1 | WP_003973134.1 | 42.0 |
| Acyltransferase | FabD | WP_016468811.1 | WP_011028322.1 | 67.4 | -- | -- | -- | -- |
| Ketosynthase | FabH | WP_016468812.1 | WP_003976418.1 | 81.9 | P | WP_016467650.1 | WP_011030514.1 | 62.9 |
| Ketoreductase | FabG[a] | WP_030408697.1 | WP_003977008.1 | 82.5 | -- | -- | -- | -- |
| Ketoreductase | FabG3[a] | WP_037612827.1 | WP_011027734.1 | 84.6 | -- | -- | -- | -- |
| Dehydratase | FabA[b][c] | WP_016473473.1; WP_016473472.1 | WP_011029784.1; WP_011029785.1 | 79.1; 81.3 | -- | -- | -- | -- |
| Enoylreductase | FabI[a] | WP_016468107.1 | WP_003977009.1 | 74.9 | -- | -- | -- | -- |
| Ketosynthase | FabF | WP_016468814.1 | WP_011028323.1 | 80.7 | R | WP_016467648.1 | WP_011030513.1 | 62.2 |
| Thioesterase | -- | -- | -- | -- | J | WP_106963665.1 | WP_003973126.1 | 51.7 |

[a] Singh and Reynolds 2015
[b] Singh and Reynolds 2016
[c] Heterodimer

**Table S5.** Pairwise analysis of primary fatty acid biosynthetic enzymes against their secondary metabolism counterparts in *S. albus* NRRL B-2362 *and S. coelicolor* A2(3).

| | *S. albus* | | | | | *S. coelicolor* | | | | |
|---|---|---|---|---|---|---|---|---|---|---|
| Function | Orthologue Name | FAS Enzyme | Orthologue Name | Tab Enzyme | BLAST %ID | Orthologue Name | FAS Enzyme | Orthologue Name | Red Enzyme | BLAST %ID |
| Acyl-carrier protein | FabC | WP_016468813.1 | Q | WP_016467649.1 | 36.8 | FabC | WP_003976417.1 | Q | WP_003973134.1 | 31.4 |
| Ketosynthase | FabH | WP_016468812.1 | P | WP_016467650.1 | 39.8 | FabH | WP_003976418.1 | P | WP_011030514.1 | 38.9 |
| Ketosynthase | FabF | WP_016468814.1 | R | WP_016467648.1 | 44.5 | FabF | WP_011028323.1 | R | WP_011030513.1 | 41.9 |

### Appendix A: Acyl S-NAC NMR and LC-MS Data

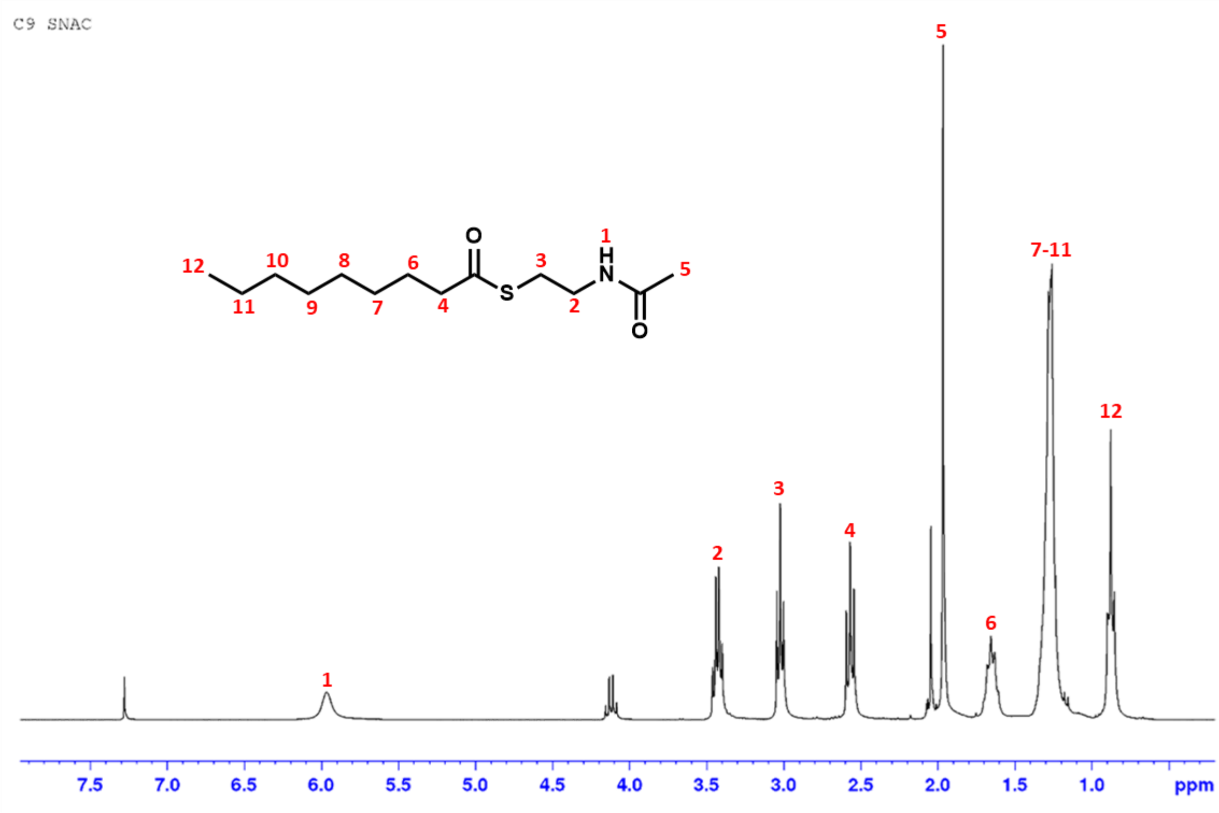


**Fig. SA1 | [1]H-NMR spectrum of C9-SNAC.** [1]H NMR (CDCl3, 300 MHz): δ 5.89 (bs, 1H), 3.43 (m, 2H), 3.02 (t, J=6.56 Hz, 2H), 2.58 (t, J=7.68 Hz, 2H), 1.97 (s, 3H), 1.66 (m, 2H), 1.35-1.27 (m, 10H), 0.89 (t, J=6.85 Hz 3H). Yield 94%.

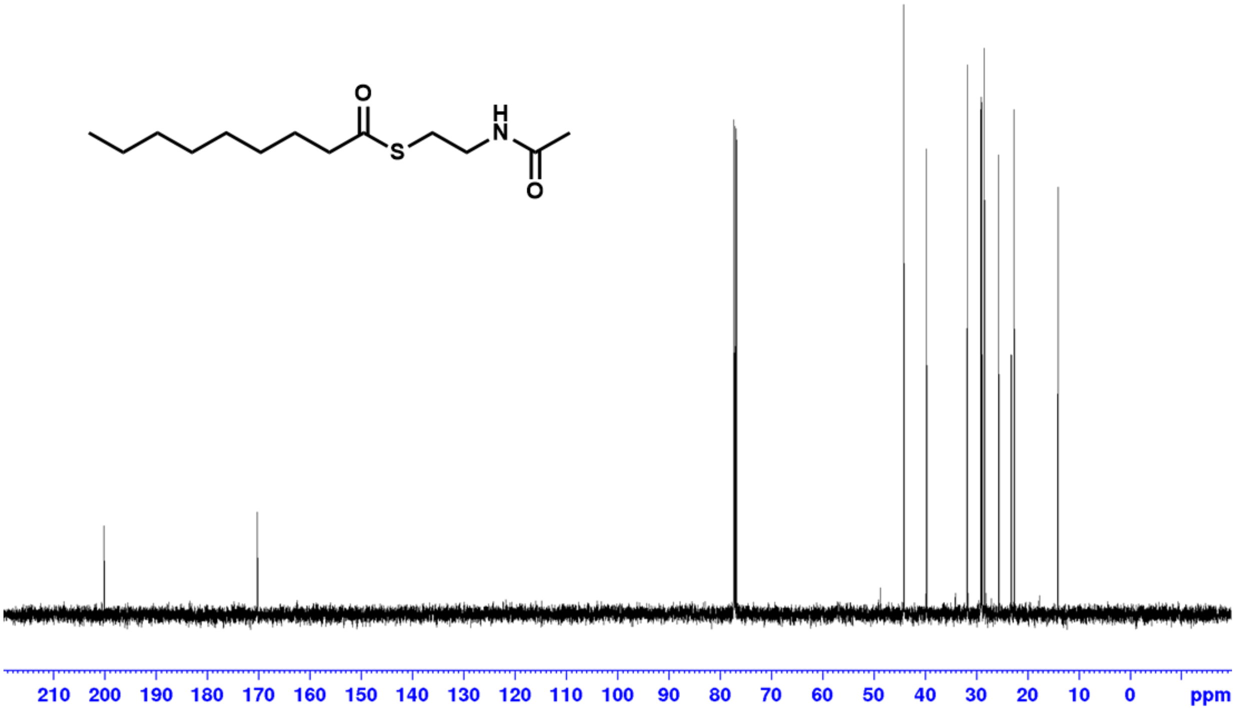


**Fig. SA2 | $^{13}$C-NMR spectrum of C$_9$-SNAC.** $^{13}$C NMR (CDCl3, 125 MHz): δ 200.2, 170.3, 44.1, 39.7, 31.8, 29.2, 29.0, 28.9, 28.4, 28.4, 25.7, 23.2, 22.6, 14.1.

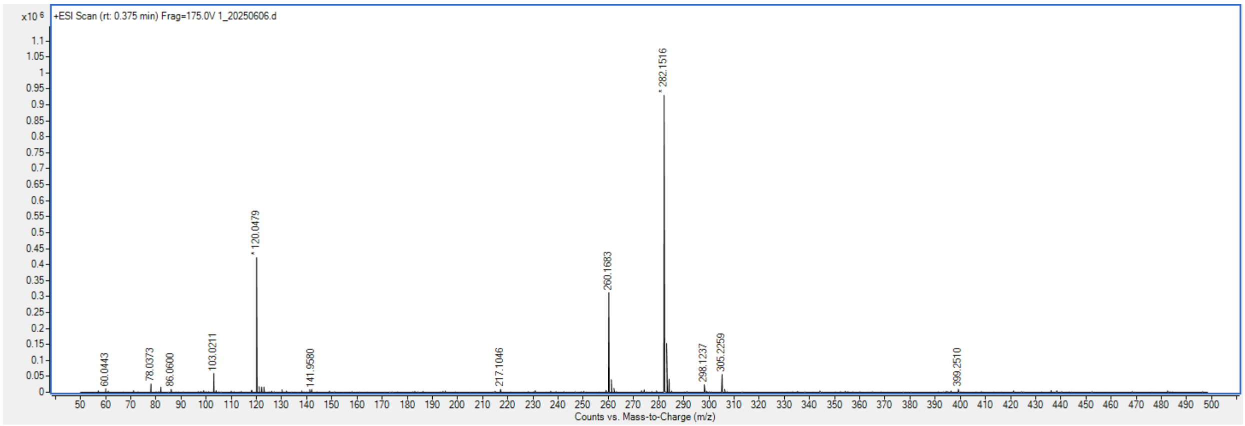


**Fig. SA3 | HR-ESI-MS of C$_9$-SNAC.** $[M+H]^+$ Calculated for $C_{13}H_{26}NO_2S$: 260.1684 m/z; found 260.1683 m/z.

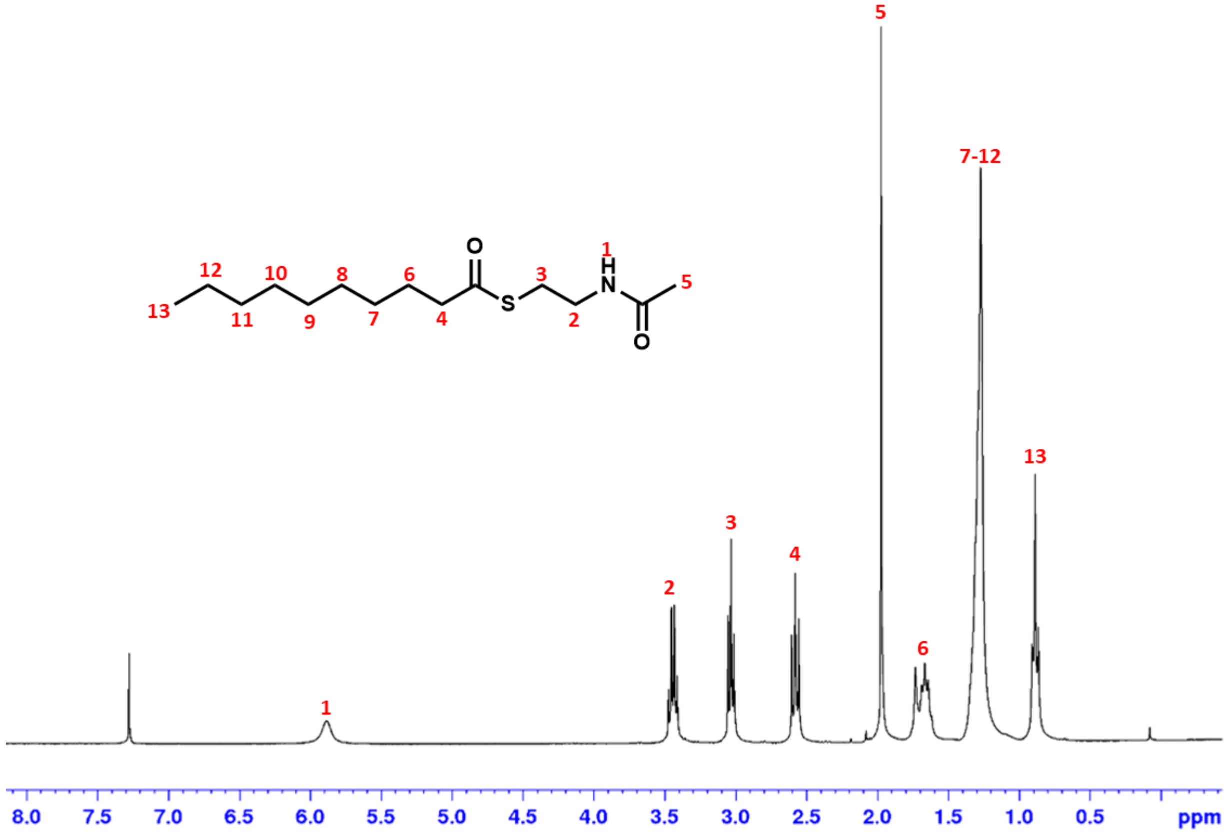


**Fig. SA4 | [1]H-NMR spectrum of $C_{10}$-SNAC.** [1]H NMR (CDCl3, 300 MHz): δ 5.89 (bs, 1H), 3.43 (m, 2H), 3.02 (t, J=6.56 Hz, 2H), 2.58 (t, J=7.68 Hz, 2H), 1.97 (s, 3H), 1.66 (m, 2H), 1.35-1.27 (m, 12H), 0.89 (t, J=6.85 Hz 3H). Yield 61%.

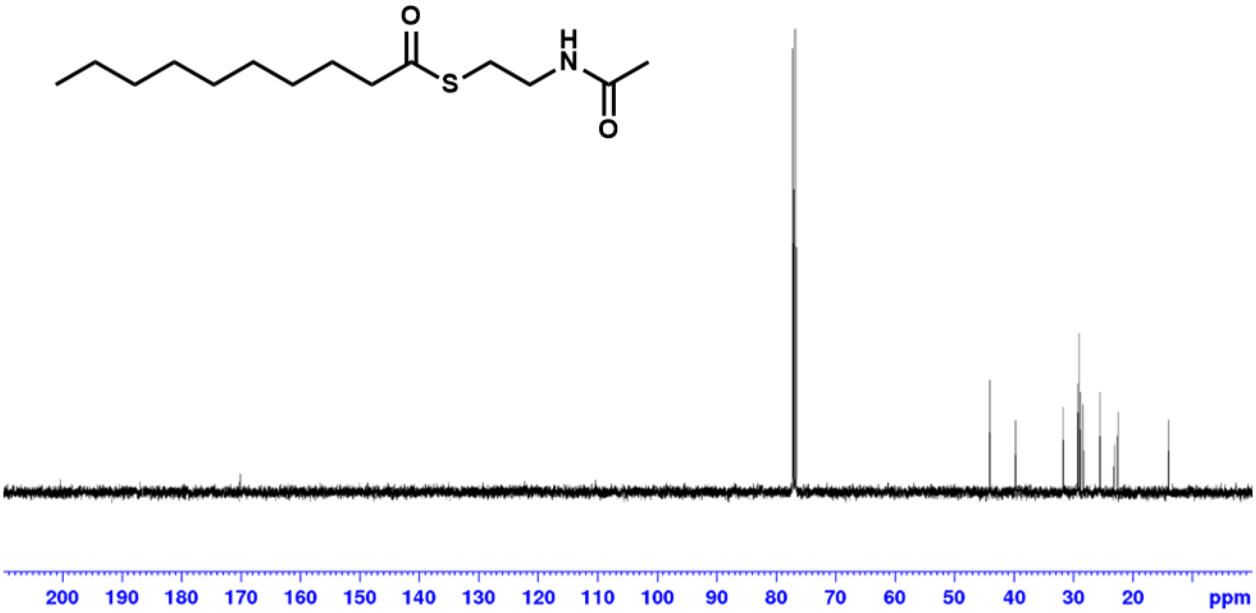


**Fig. SA5 | $^{13}$C-NMR spectrum of $C_{10}$-SNAC.** $^{13}$C NMR (CDCl3, 125 MHz): δ 200.3, 170.2, 44.1, 39.8, 31.9, 29.4, 29.2, 28.9, 28.4, 25.7, 23.2, 22.7, 14.1.

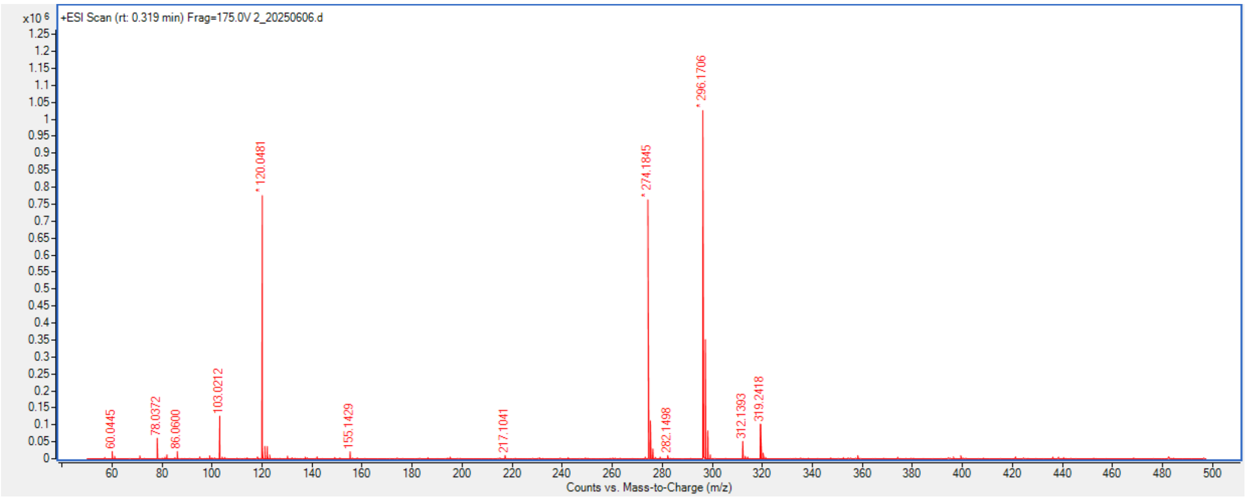


**Fig. SA6 | HR-ESI-MS of $C_{10}$-SNAC.** $[M+H]^+$ Calculated for $C_{14}H_{28}NO_2S$: 274.1840 m/z; found 274.1645 m/z.

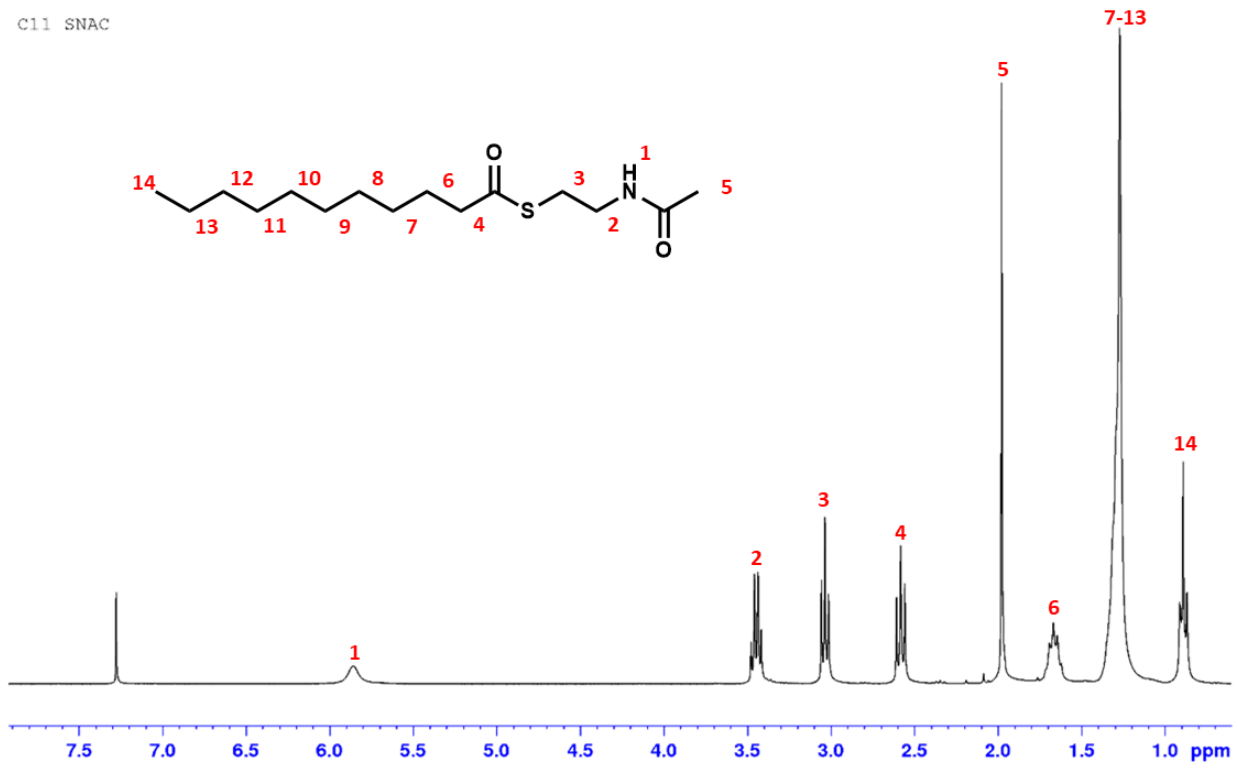


**Fig. SA7 | [1]H-NMR spectrum of C11-SNAC.** [1]H NMR (CDCl3, 300 MHz): δ 5.86 (bs, 1H), 3.44 (m, 2H), 3.03 (t, J=6.56 Hz, 2H), 2.58 (t, J=7.65 Hz, 2H), 1.97 (s, 3H), 1.66 (m, 2H), 1.35-1.27 (m, 14H), 0.89 (t, J=6.89 Hz 3H). Yield 45%.

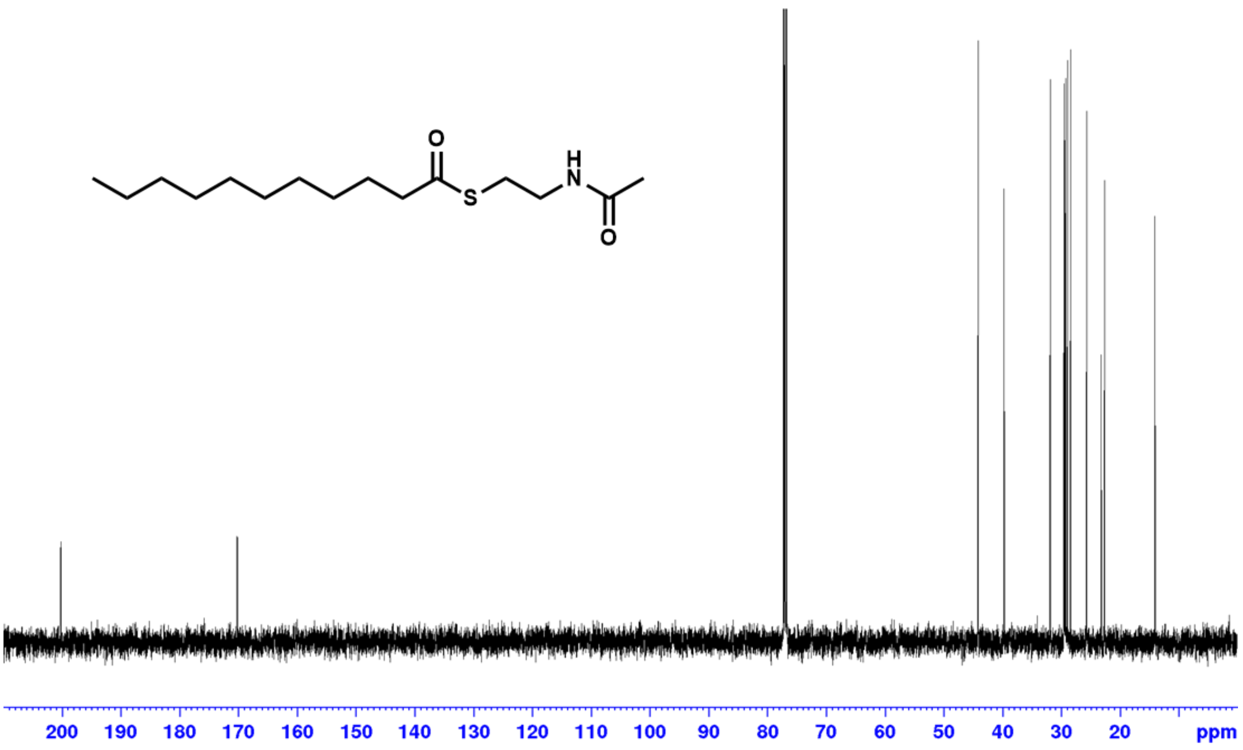

**Fig. SA8 | $^{13}$C-NMR spectrum of $C_{11}$-SNAC.** $^{13}$C NMR (CDCl3, 125 MHz): δ 200.3, 170.2, 44.1, 39.8, 31.9, 29.6-29.5, 29.4, 29.3 29.2, 28.9, 28.4, 25.7, 23.2, 22.7, 14.1.

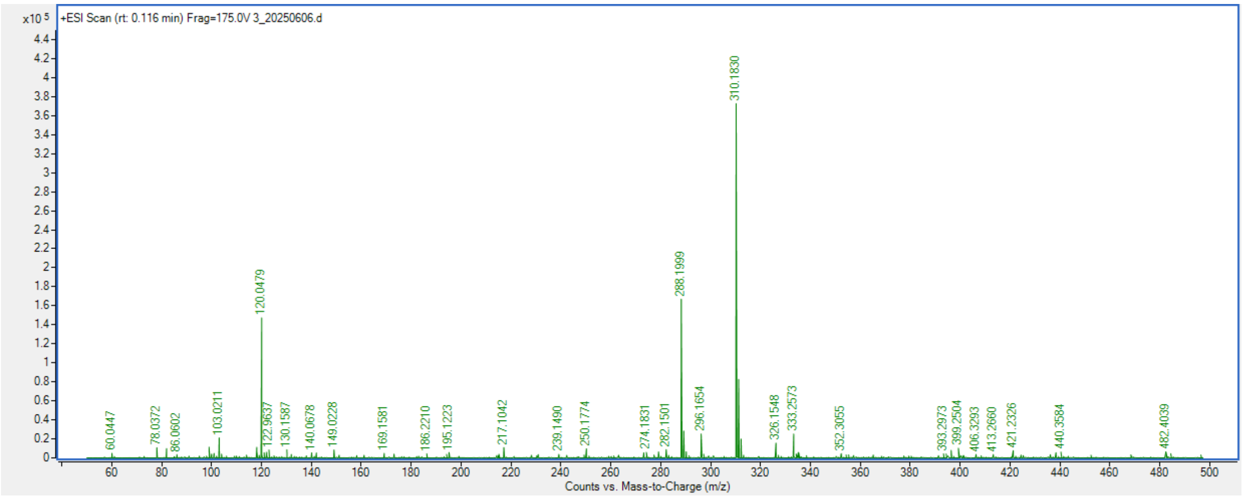


**Fig. SA9 | HR-ESI-MS of $C_{11}$-SNAC.** $[M+H]^+$ Calculated for $C_{15}H_{30}NO_2S$: 288.1997 m/z; found 288.1999 m/z.

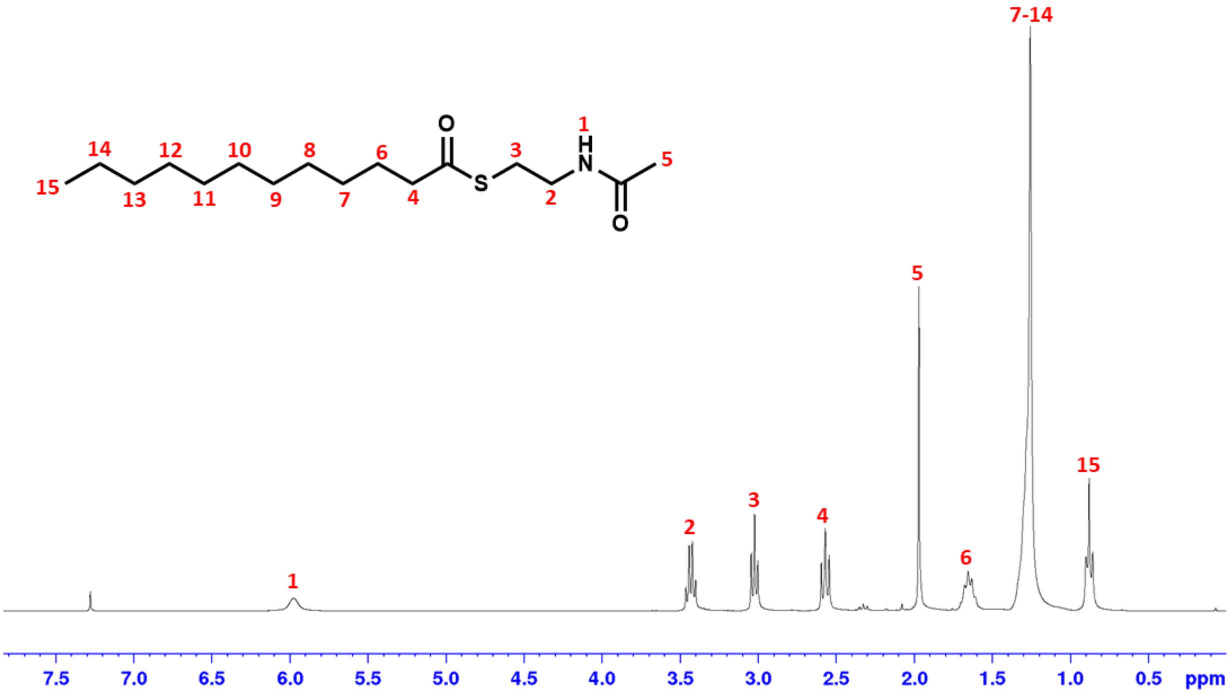


**Fig. SA10 | $^{1}$H-NMR spectrum of $C_{12}$-SNAC.** $^{1}$H NMR (CDCl3, 300 MHz): δ 5.98 (bs, 1H), 3.43 (m, 2H), 3.02 (t, J=6.41 Hz, 2H), 2.57 (t, J=7.39 Hz, 2H), 1.97 (s, 3H), 1.65 (m, 2H), 1.33-1.26 (m, 16H), 0.88 (t, J=6.33 Hz 3H). Yield 47%.

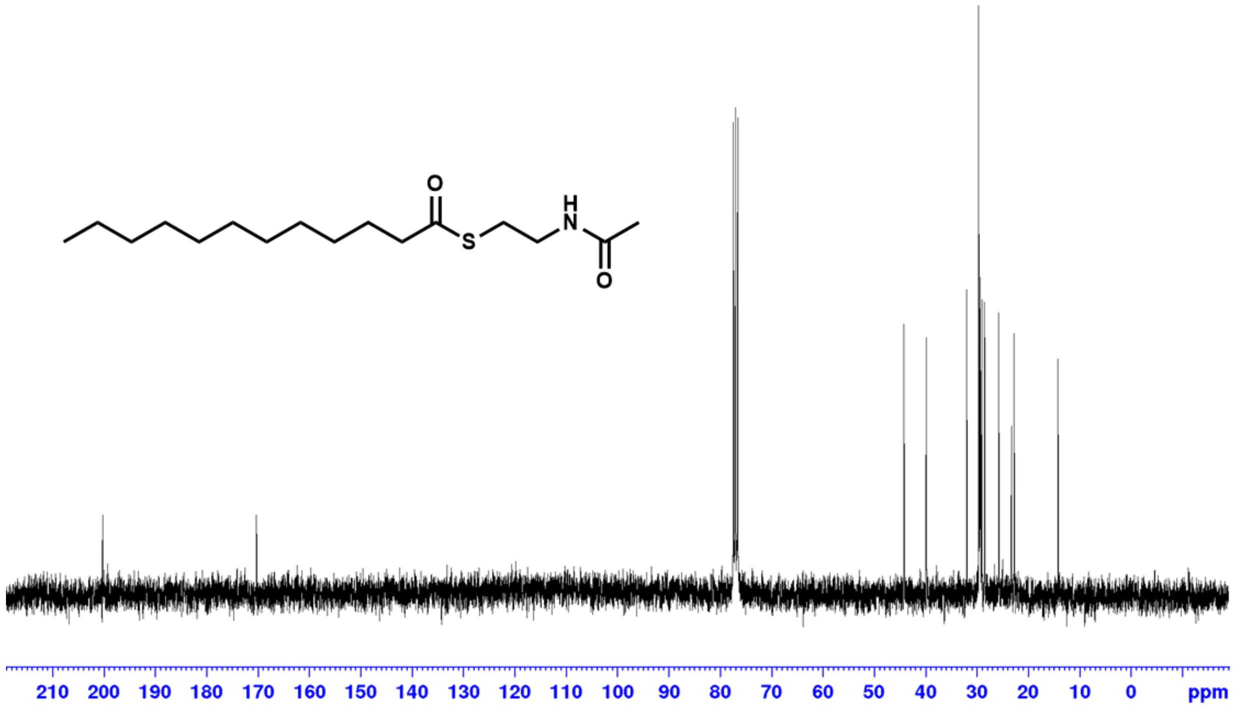


**Fig. SA11 | $^{13}$C-NMR spectrum of $C_{12}$-SNAC.** $^{13}$C NMR (CDCl3, 75 MHz): δ 200.3, 170.4, 44.1, 39.8, 31.9, 29.6-29.5, 29.4, 29.3, 29.2, 28.9, 28.4, 25.7, 23.1, 22.7, 14.1.

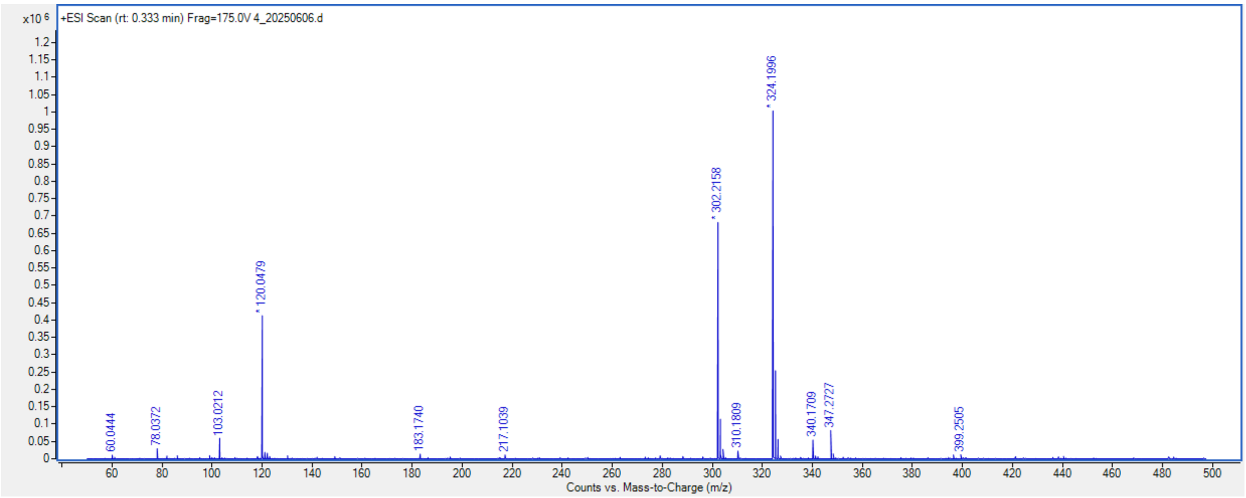


**Fig. SA12 | HR-ESI-MS of $C_{12}$-SNAC.** $[M+H]^+$ Calculated for $C_{16}H_{32}NO_2S$: 302.2154 m/z; found 302.2158 m/z.

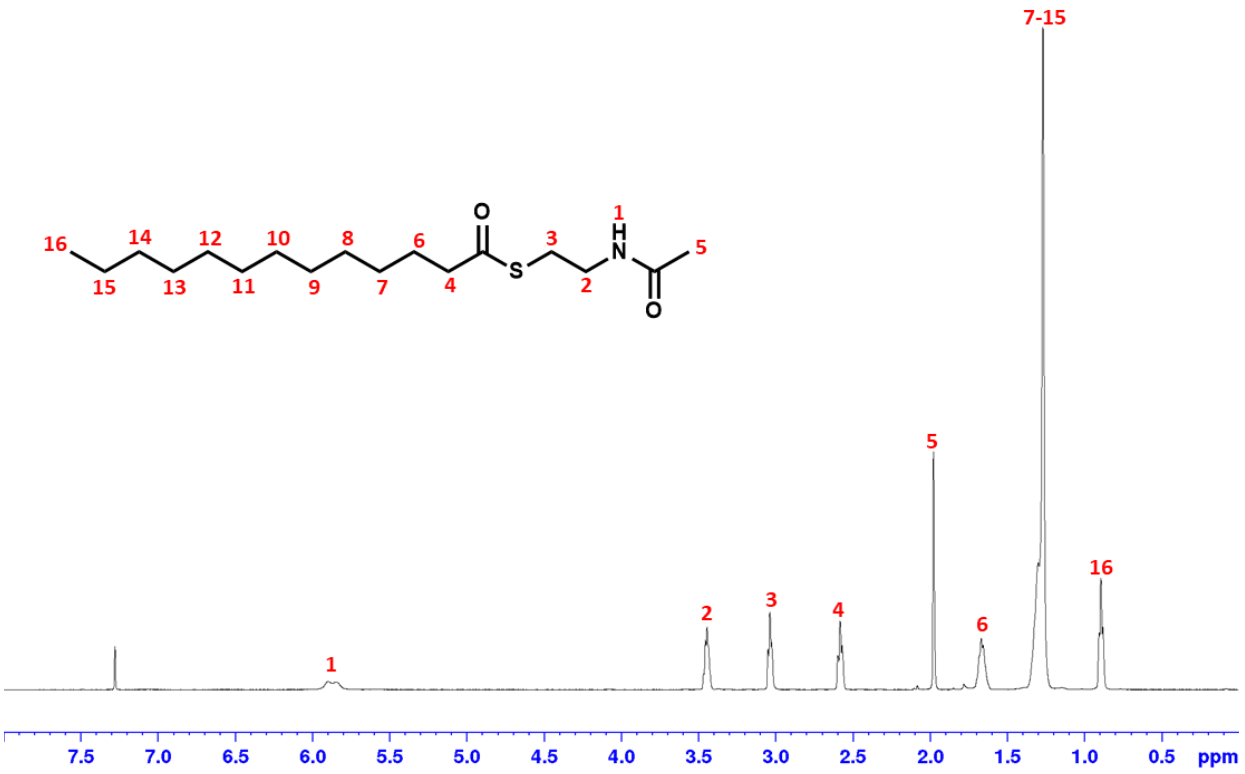


**Fig. SA13 | ¹H-NMR spectrum of $C_{13}$-SNAC.** ¹H NMR (CDCl3, 500 MHz): δ 5.87 (bs, 1H), 3.44 (m, 2H), 3.04 (t, J=5.27 Hz, 2H), 2.58 (t, J=7.28 Hz, 2H), 1.98 (s, 3H), 1.67 (m, 2H), 1.34-1.26 (m, 18H), 0.88 (t, J=5.99 Hz 3H). Yield 78%.

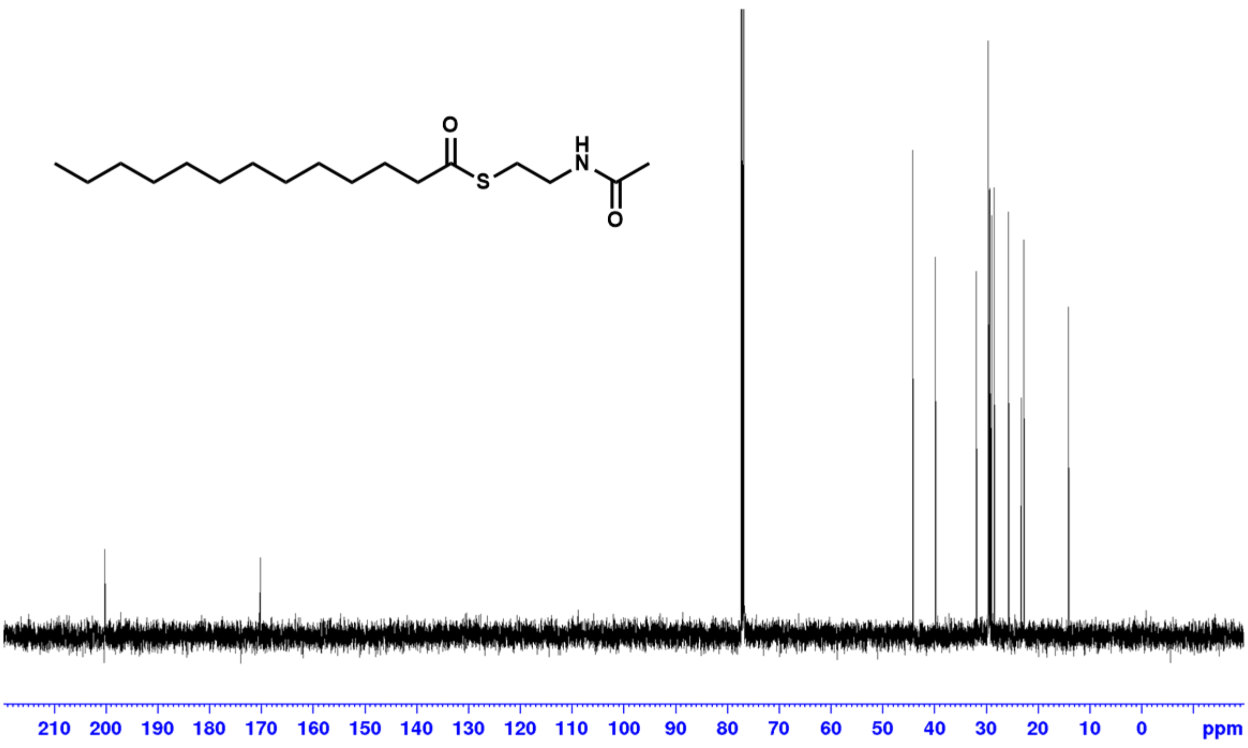

**Fig. SA14 | $^{13}$C-NMR spectrum of $C_{13}$-SNAC.** $^{13}$C NMR (CDCl3, 125 MHz): δ 200.3, 170.2, 44.1, 39.8, 31.9, 29.7-29.6, 29.4, 29.3, 29.2, 28.9, 28.4, 25.7, 23.2, 22.7, 14.1.

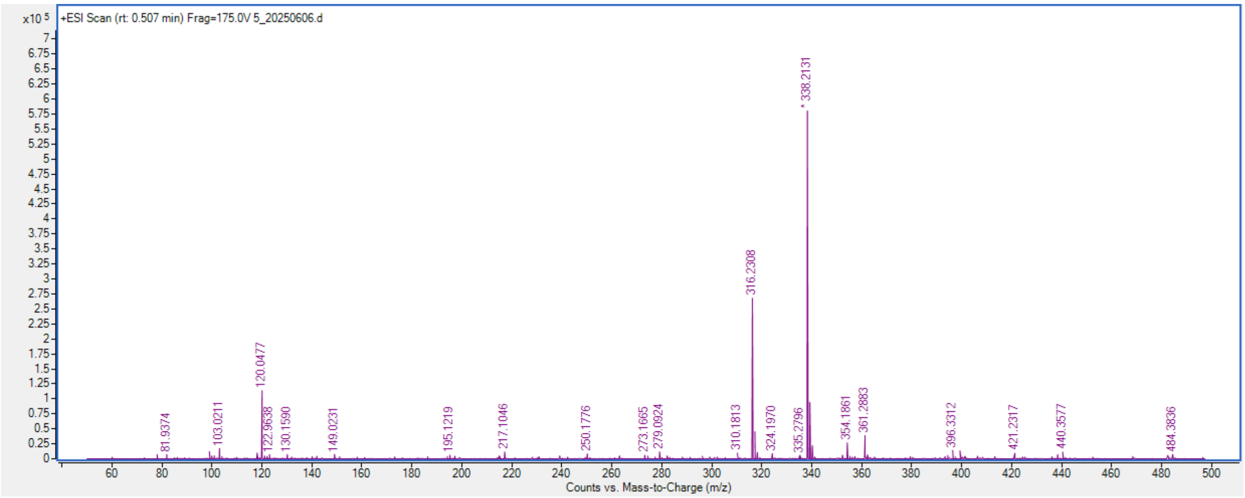


**Fig. SA15 | HR-ESI-MS of $C_{13}$-SNAC.** $[M+H]^+$ Calculated for $C_{17}H_{34}NO_2S$: 316.2310 m/z; found 316.2308 m/z.

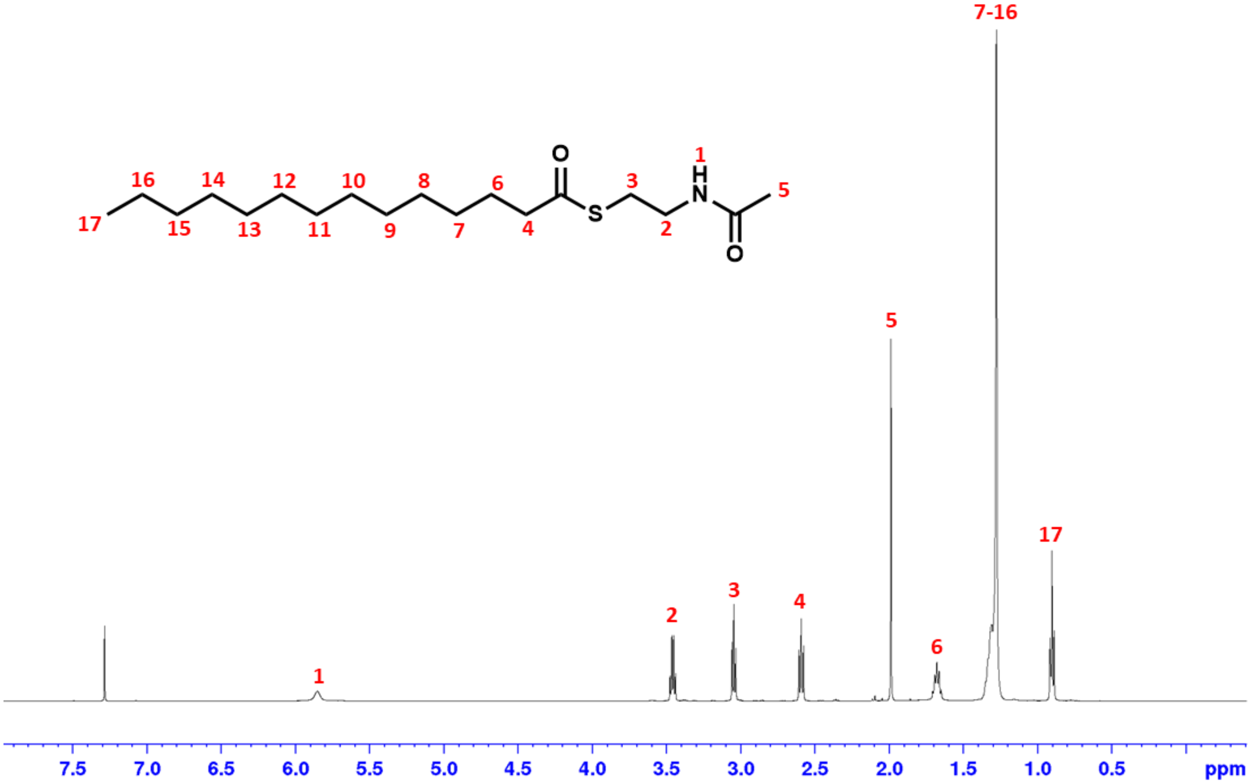

**Fig. SA16 | $^{1}$H-NMR spectrum of $C_{14}$-SNAC.** $^{1}$H NMR (CDCl3, 500 MHz): δ 5.85 (bs, 1H), 3.46 (m, 2H), 3.04 (t, J=6.59 Hz, 2H), 2.59 (t, J=7.65 Hz, 2H), 1.99 (s, 3H), 1.68 (m, 2H), 1.34-1.28 (m, 16H), 0.90 (t, J=6.90 Hz 3H). Yield 42%.

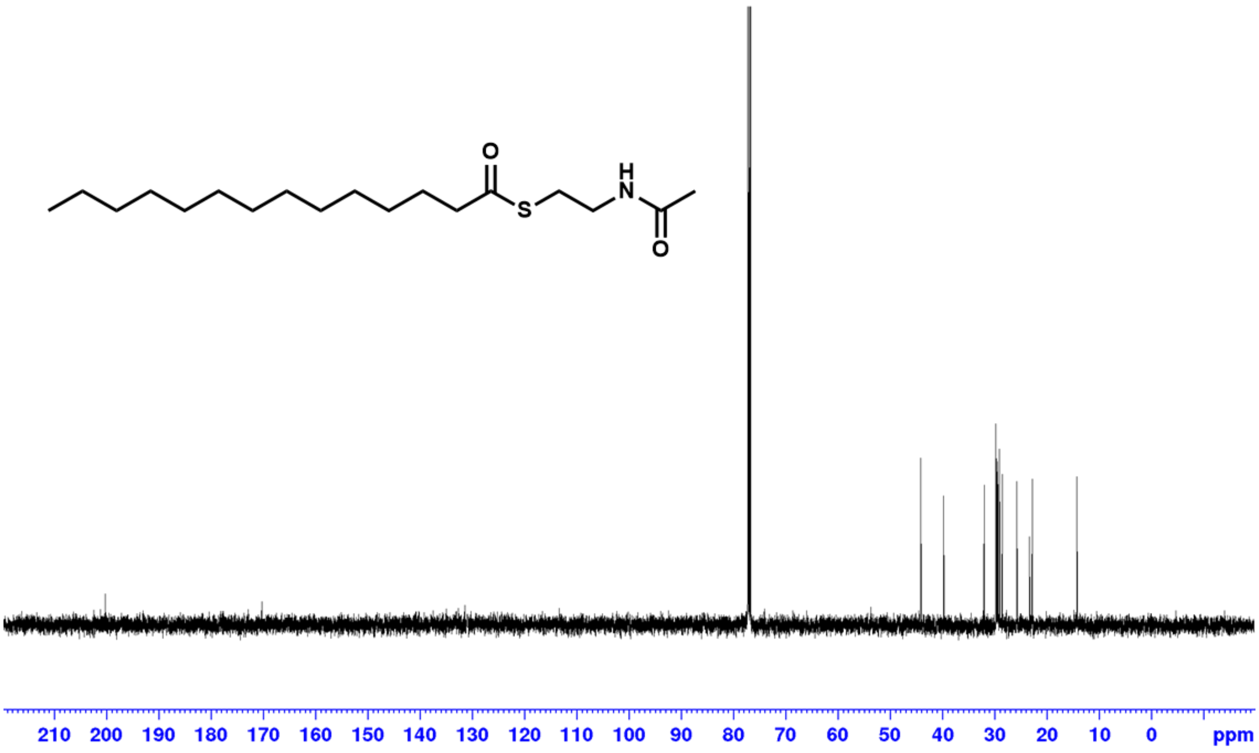


**Fig. SA17 | $^{13}$C-NMR spectrum of $C_{14}$-SNAC.** $^{13}$C NMR (CDCl3, 125 MHz): δ 200.3, 170.2, 44.1, 39.8, 31.9, 29.7-29.6, 29.4, 29.3, 29.2, 28.9, 28.4, 25.7, 23.2, 22.7, 14.1.

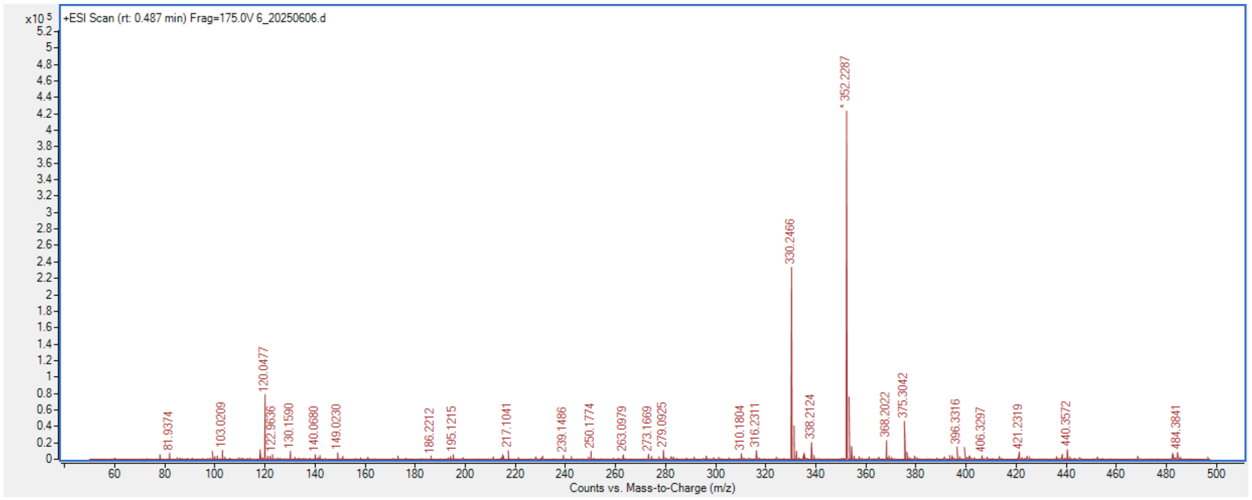


**Fig. SA18 | HR-ESI-MS of $C_{14}$-SNAC.** $[M+H]^{+}$ Calculated for $C_{18}H_{36}NO_2S$: 330.2467 m/z; found 330.2466 m/z.

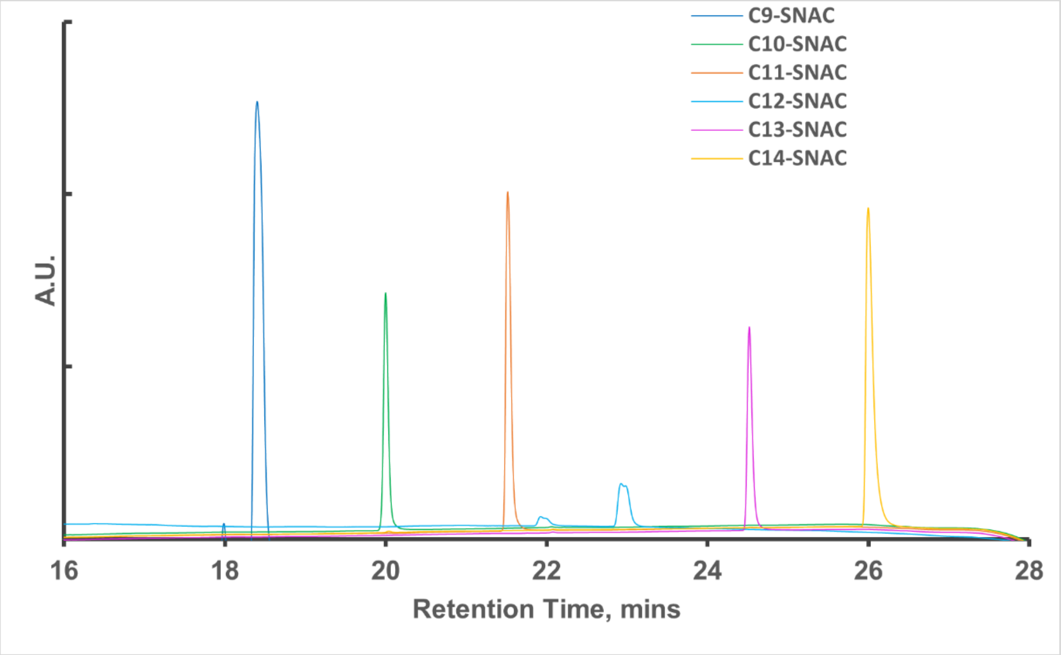


**Fig. SA19 | UPLC photodiode array traces of purified acyl-SNACs.**

### Appendix B: Plasmid Maps

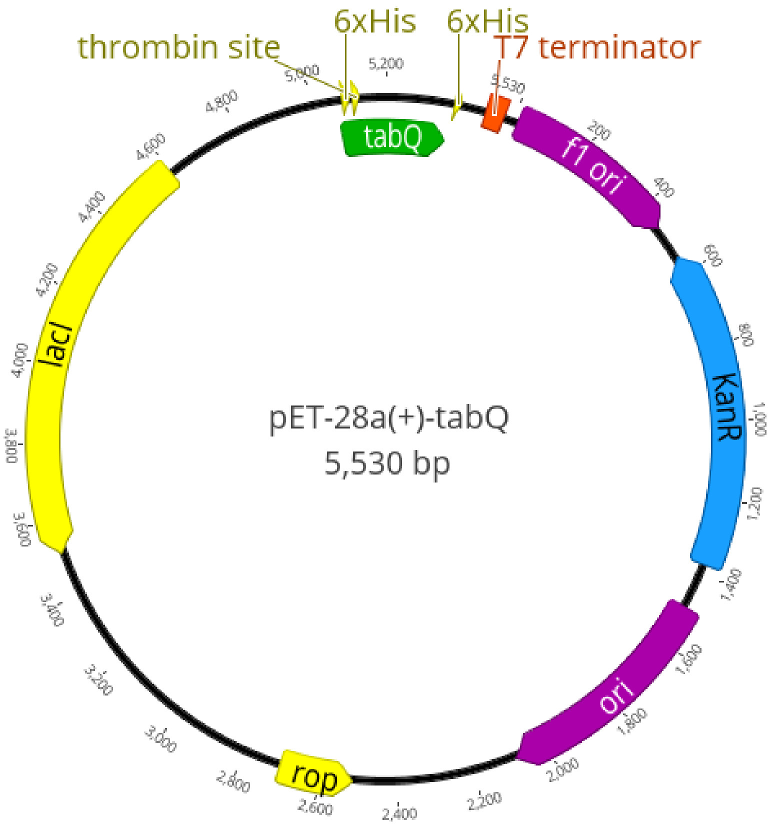


**Fig. SB1 | Plasmid map and nucleotide sequence of pET-28a-tabQ.** Sequence of tabQ shown in red.

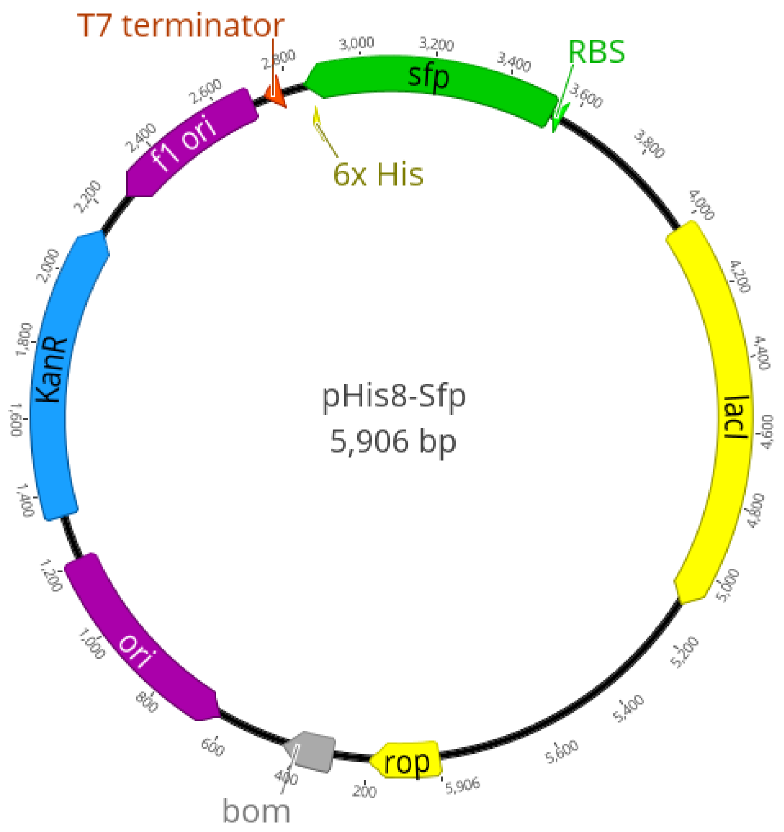


**Fig. SB2 | Plasmid map and nucleotide sequence of pHIS8-Sfp.** Sequence of Sfp shown in red.

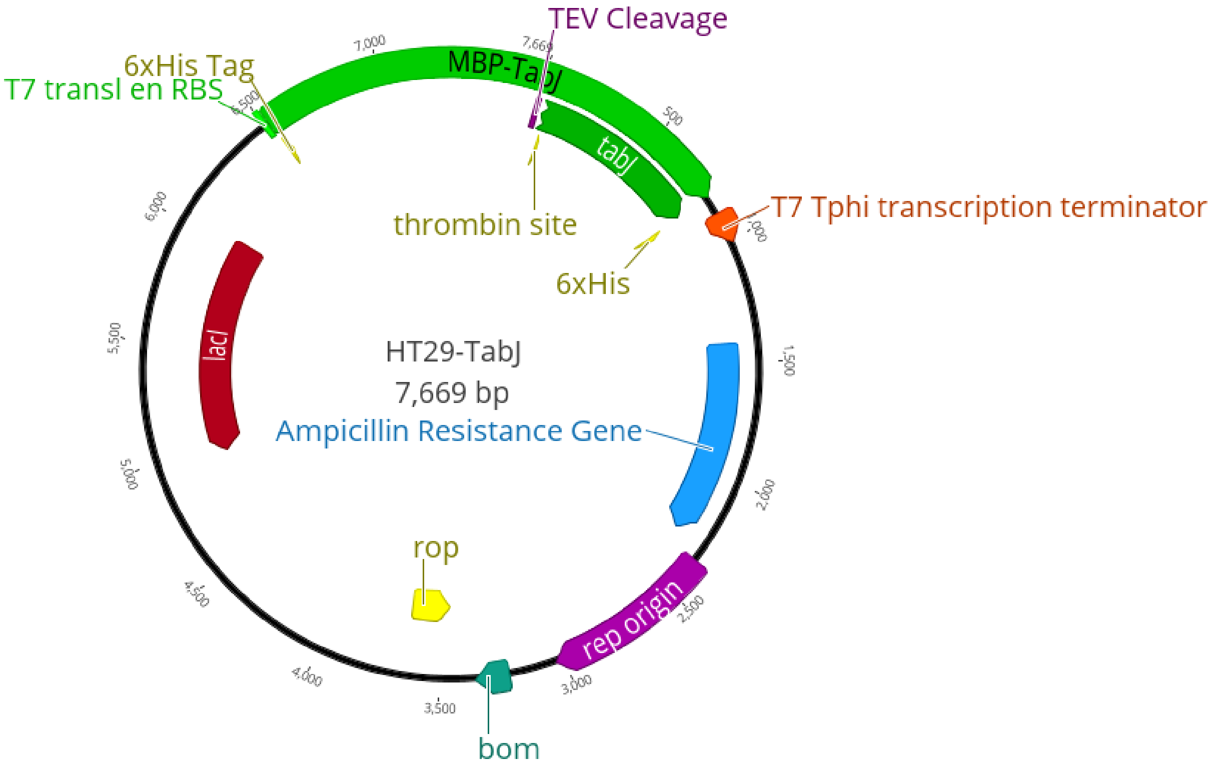


**Fig. SB3 | Plasmid map and nucleotide sequence of HT29-tabJ.** Sequence of tabJ shown in red and the sequence of the maltose-binding protein is shown in blue.

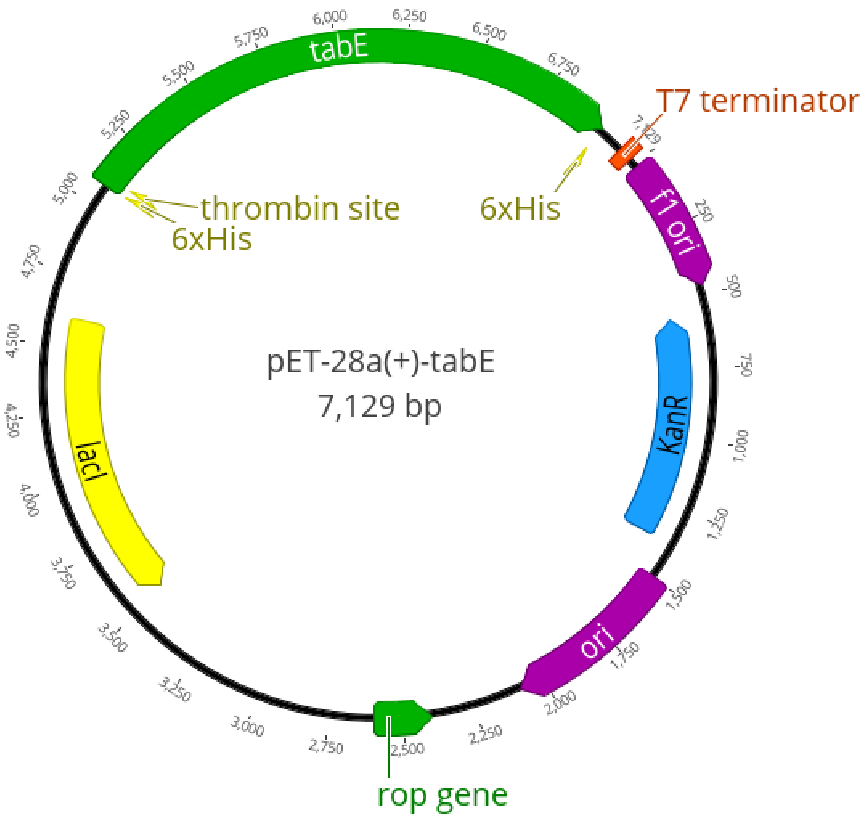


**Fig. SB4 | Plasmid map and nucleotide sequence of pET-28a-tabE.** Sequence of tabE shown in red.

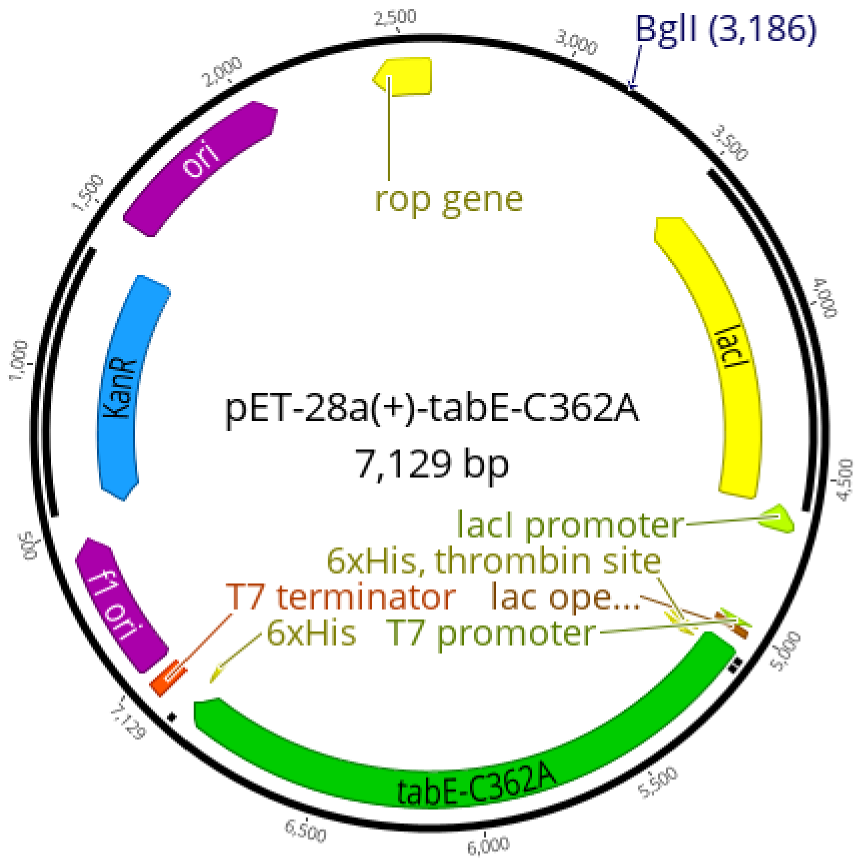


**Fig. SB5 | Plasmid map and nucleotide sequence of pET-28a-tabE-C362A.** Sequence of tabE-C362A shown in red.

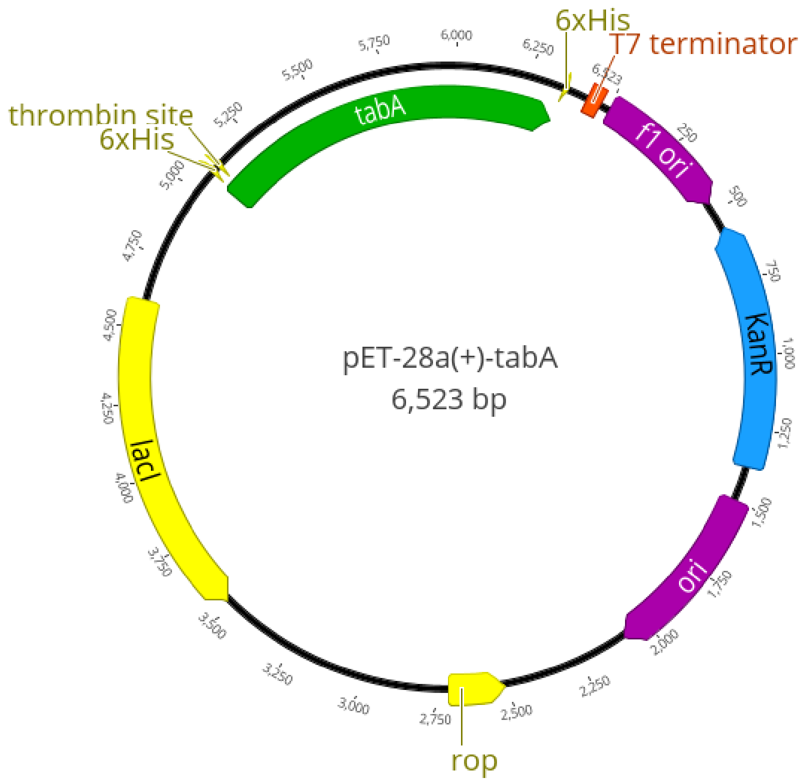


**Fig. SB6 | Plasmid map and nucleotide sequence of pET-28a-tabA.** Sequence of tabA shown in red.

### Appendix C: TabJ $C_{12}$-TabQ Assay Raw Data

| Time, h | Average Abs No Enzyme (n=2) | Average Abs TabJ+C12-TabQ (n=3) |
|---|---|---|
| 0.033333 | 0.3295 | 0.315 |
| 0.066667 | 0.335 | 0.318 |
| 0.1 | 0.3395 | 0.318666667 |
| 0.133333 | 0.3445 | 0.320333333 |
| 0.166667 | 0.3505 | 0.322 |
| 0.2 | 0.3515 | 0.323666667 |
| 0.233333 | 0.356 | 0.325333333 |
| 0.266667 | 0.356 | 0.327 |
| 0.3 | 0.3585 | 0.328666667 |
| 0.333333 | 0.3615 | 0.329666667 |
| 0.366667 | 0.3585 | 0.332 |
| 0.4 | 0.3565 | 0.333666667 |
| 0.433333 | 0.358 | 0.334666667 |
| 0.466667 | 0.3575 | 0.337 |
| 0.5 | 0.355 | 0.336666667 |
| 0.533333 | 0.353 | 0.337666667 |
| 0.566667 | 0.354 | 0.337666667 |
| 0.6 | 0.3555 | 0.338333333 |
| 0.633333 | 0.356 | 0.339333333 |
| 0.666667 | 0.35 | 0.339333333 |
| 0.7 | 0.358 | 0.339666667 |
| 0.733333 | 0.3495 | 0.340333333 |
| 0.766667 | 0.348 | 0.340333333 |
| 0.8 | 0.3465 | 0.340666667 |
| 0.833333 | 0.3465 | 0.340666667 |
| 0.866667 | 0.3445 | 0.340666667 |
| 0.9 | 0.344 | 0.340666667 |
| 0.933333 | 0.3445 | 0.341333333 |
| 0.966667 | 0.3435 | 0.341 |
| 1 | 0.3415 | 0.341 |
| 1.033333 | 0.3415 | 0.341333333 |
| 2.033333 | 0.331 | 0.341666667 |
| 3.033333 | 0.331 | 0.342666667 |
| 4.033333 | 0.3295 | 0.344666667 |
| 5.033333 | 0.3275 | 0.346 |
| 6.033333 | 0.3265 | 0.347 |
| 7.033333 | 0.324 | 0.348 |
| 8.033333 | 0.3225 | 0.347666667 |

| | | |
|---|---|---|
| 9.033333 | 0.32 | 0.348 |
| 10.03333 | 0.3185 | 0.348333333 |
| 11.03333 | 0.3175 | 0.348333333 |
| 12.03333 | 0.315 | 0.348333333 |
| 13.03333 | 0.3135 | 0.348 |
| 14.03333 | 0.312 | 0.348333333 |
| 15.03333 | 0.3105 | 0.348 |
| 16.03333 | 0.309 | 0.348333333 |
| 17.03333 | 0.3085 | 0.349 |
| 18.03333 | 0.307 | 0.348666667 |
| 19.03333 | 0.306 | 0.349333333 |
| 20.03333 | 0.305 | 0.35 |
| 21.03333 | 0.3035 | 0.35 |
| 22.03333 | 0.3025 | 0.351333333 |
| 23.03333 | 0.301 | 0.352 |
| 24.03333 | 0.3 | 0.352666667 |
| 25.03333 | 0.299 | 0.353666667 |
| 26.03333 | 0.2985 | 0.356 |

### Appendix D: Protein Purification SDS-PAGE

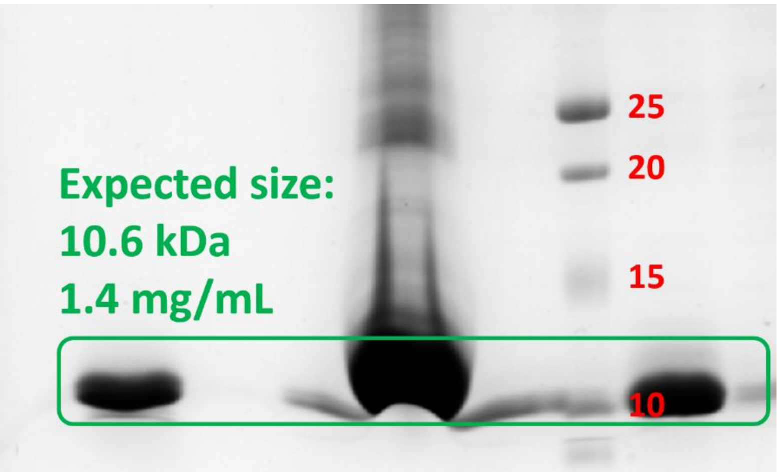


**Fig. SD1 | Nickel affinity chromatography purification of TabQ.**

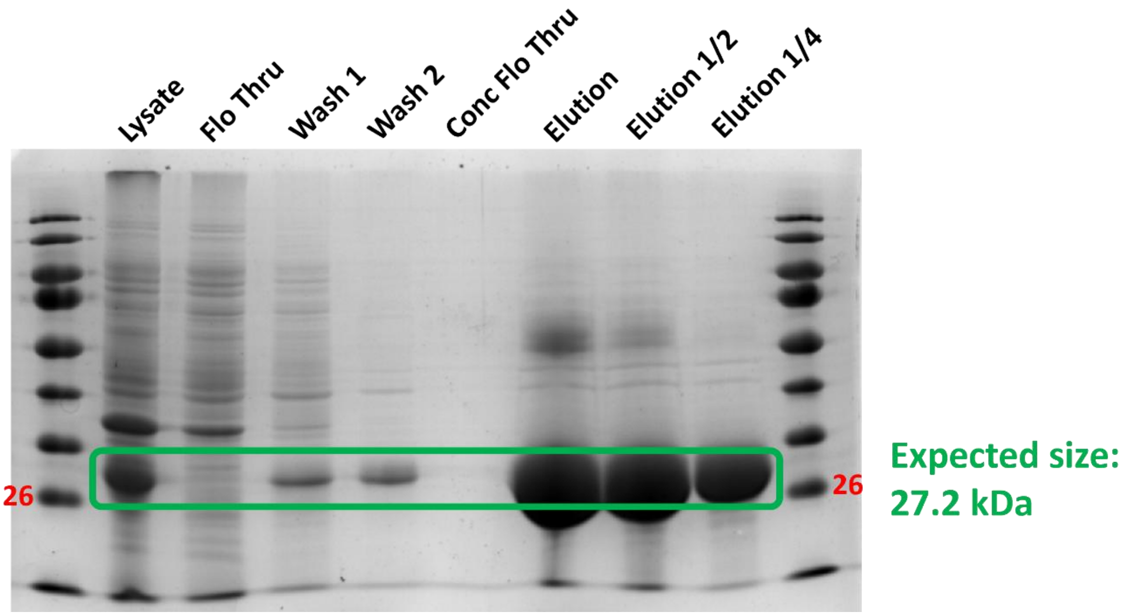


**Fig. SD2 | Nickel affinity chromatography purification of Sfp.**

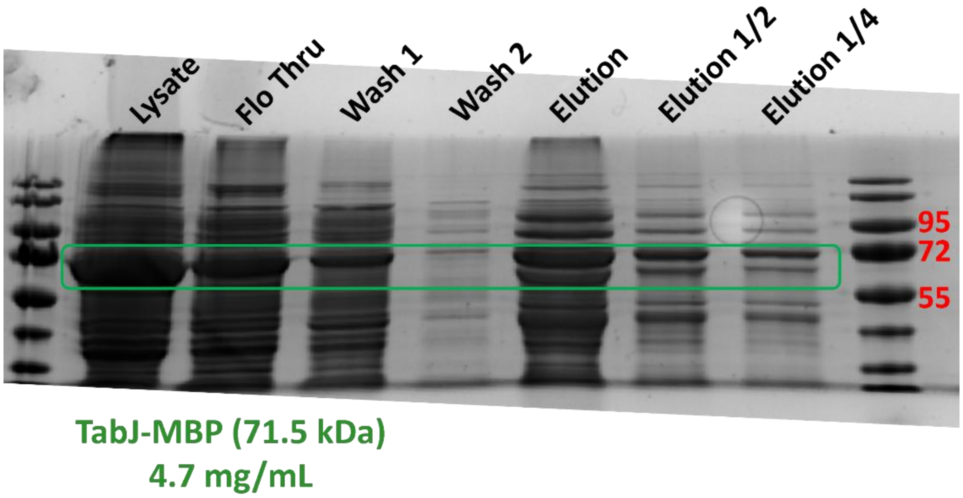


**Fig. SD3 | Nickel affinity chromatography purification of TabJ-MBP.**

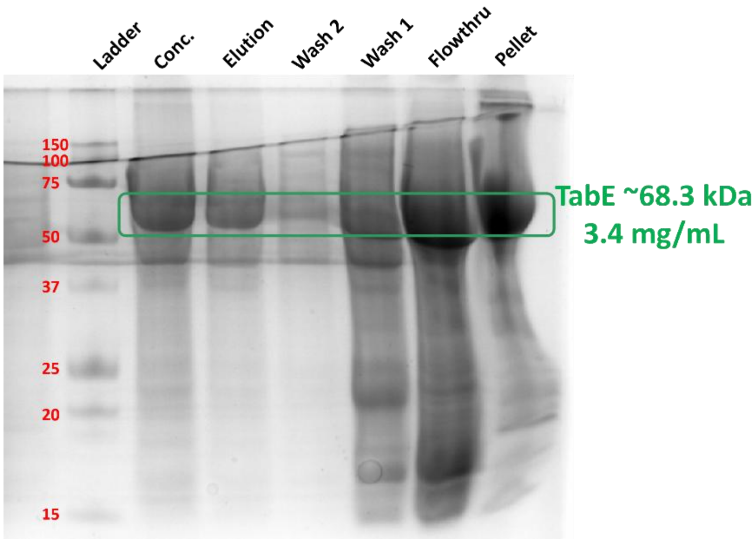


**Fig. SD4 | Nickel affinity chromatography purification of TabE.**

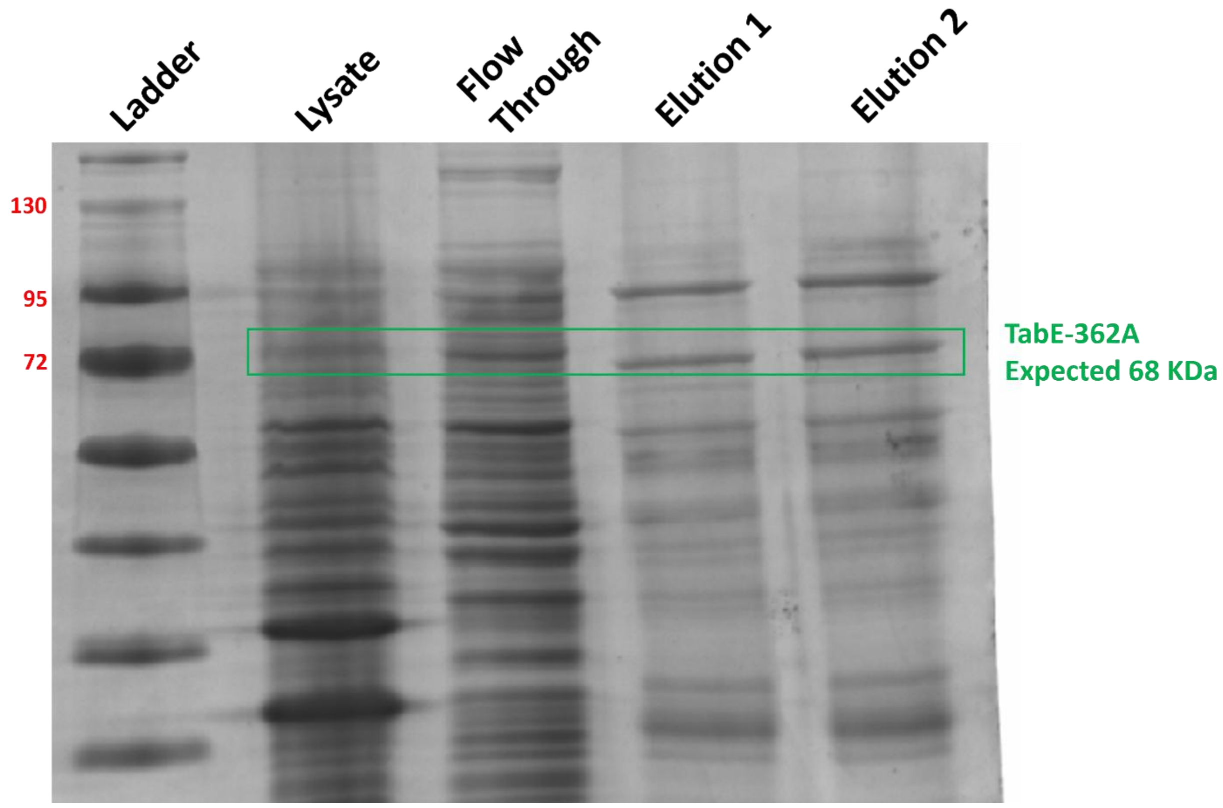


**Fig. SD5 | Nickel affinity chromatography purification of TabE-C362A.**

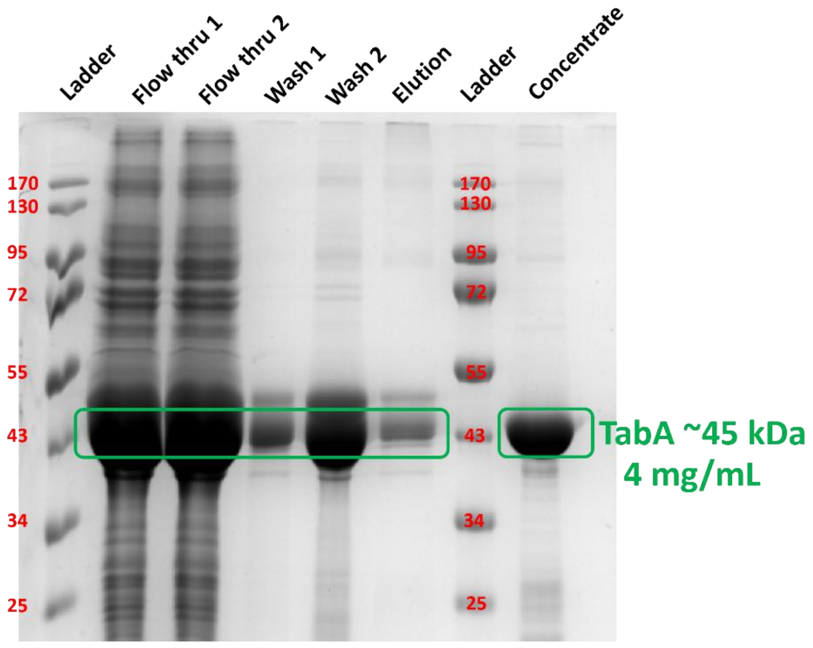


**Fig. SD6 | Nickel affinity chromatography purification of TabA.**
